# A Putative Binding Model of Nitazene Derivatives at the *μ*-Opioid Receptor

**DOI:** 10.1101/2024.10.03.616560

**Authors:** Joseph Clayton, Lei Shi, Michael J. Robertson, Georgios Skiniotis, Michael Michaelides, Lidiya Stavitskaya, Jana Shen

## Abstract

Nitazenes are a class of novel synthetic opioids with exceptionally high potency. Currently, an experimental structure of *µ*OR-opioid receptor (*µ*OR) in complex with a nitazene is lacking. Here we used a suite of computational tools, including consensus docking, conventional molecular dynamics (MD) and metadynamics simulations, to investigate the *µ*OR binding modes of nitro-containing meto-, eto-, proto-, buto-, and isotonitazenes and nitro-less analogs, metodes-, etodes-, and protodesnitazenes. Docking generated three binding modes, whereby the nitro-substituted or unsubstituted benzimidazole group extends into SP1 (subpocket 1 between transmembrane helix or TM 2 and 3), SP2 (subpocket 2 between TM1, TM2, and TM7) or SP3 (subpocket 3 between TM5 and TM6). Simulations suggest that etonitazene and likely also other nitazenes favor the SP2-binding mode. Comparison to the experimental structures of *µ*OR in complex with BU72, fentanyl, and mitragynine pseudoindoxyl (MP) allows us to propose a putative model for *µ*OR-ligand recognition in which ligand can access hydrophobic SP1 or hydrophilic SP2, mediated by the conformational change of Gln124^2.60^. Interestingly, in addition to water-mediated hydrogen bonds, the nitro group in nitazenes forms a *π*-hole interaction with the conserved Tyr75^1.39^. Our computational analysis provides new insights into the mechanism of *µ*OR-opioid recognition, paving the way for investigations of the structure-activity relationships of nitazenes.

**TOC Graphic:** 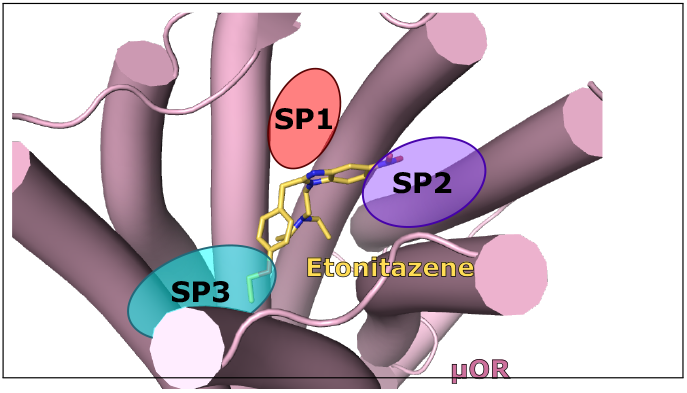

## Introduction

The number of deaths due to illicit and prescription opioid overdose has been sharply increasing in the United States, with over 80,000 deaths reported in 2022. Furthermore, over 80% of the deaths are due to synthetic opioids, primarily fentanyl.^1^ Fentanyl and other opioids are agonists of the *µ*-opioid receptor (*µ*OR), a member of the class A family of G-protein coupled receptors (GPCRs). When bound to an agonist, *µ*OR undergoes a conformational change that allows it to associate with G-proteins and *β*-arrestins; it is hypothesized that the G-protein signalling pathway produces an analgesic effect while the *β*-arrestin pathway leads to adverse effects, including respiratory depression.^2,3^ Two antagonists at the *µ*OR, naloxone and nalmefene, have been approved by the FDA as rescue agents to rapidly reverse opioid overdose through competitive binding with agonists such as fentanyl.

In recent years, a new class of synthetic opioids from the 2-benzylbenzimidazole family (also called nitazenes) has emerged on the illicit market (Figure 1). Many of the nitazene derivatives were first invented by a Swiss company in the late 1950s, but were never marketed due to their high potency.^4^ Isotonitazene was first found in biological samples in the United States in 2019 and was the subject of a public health warning from the DEA Washington Division in 2022;^5^ since then, ten nitazenes have been scheduled by the DEA.^6^ Members of this family are known to be up to 20 times more potent than fentanyl.^7–9^ A recent study found that overdose patients in the emergency department testing positive for nitazenes required a higher dose of naloxone compared to fentanyl alone and that metonitazene overdose was associated with cardiac arrest and death at a higher rate than other substances. ^10^

**Figure 1:**
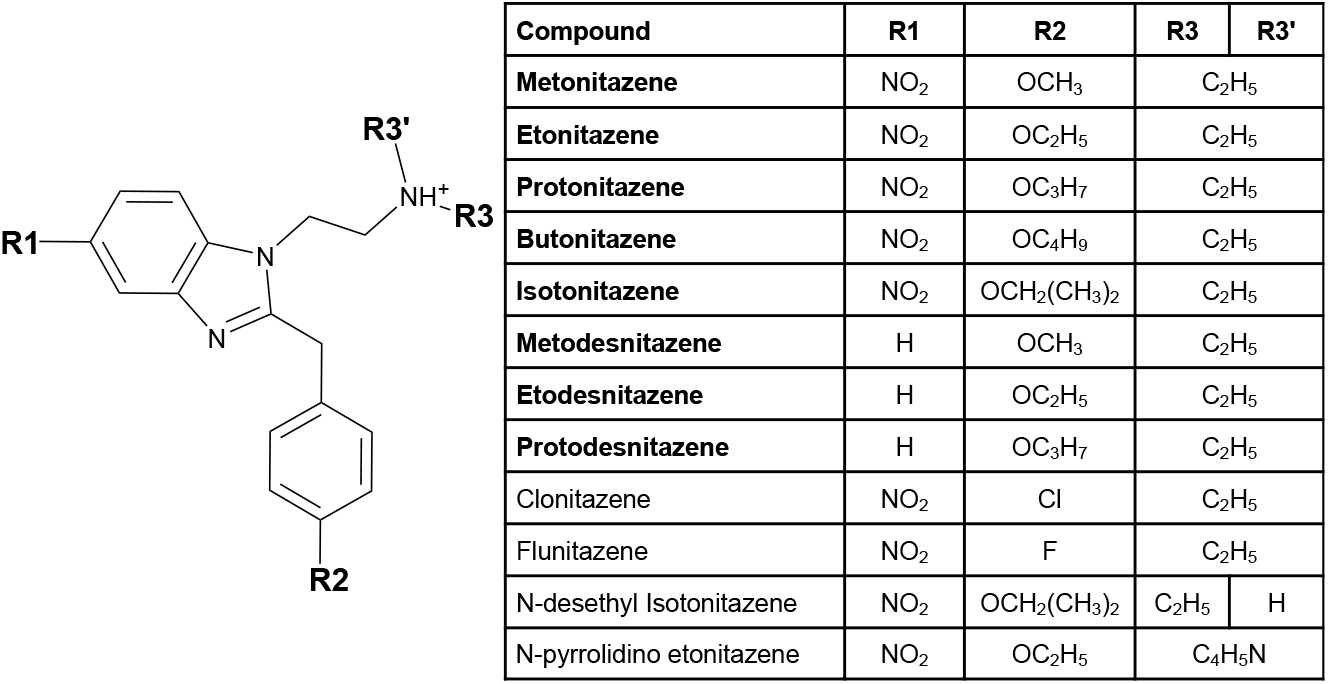
Chemical structures of nitazenes. Nitazenes share a 2-benzylbenzimidazol core with varying substitutions at R_1_, R_2_ and R_3_ positions. R_1_ is either occupied by a nitro (NO_2_) group or is empty; R_2_ is occupied by either a halogen or an - oxy group of varying sizes; R_3_ and R_3_’ are most commonly occupied by ethyl groups, but can also be part of a ring structure such as pyrrolidine. The nitazenes studied in this work are in bold font; all except protodesnitazene are currently Schedule 1 substances.

Figure 1 shows the chemical structure of selected nitazenes. Recent experimental studies^7,8^ showed that nitazenes that contain both a nitro group at R_1_ and an alkoxy group at R_2_ have a higher potency than other nitazenes. For example, etonitazene (R_1_= NO_2_, R_2_=OC_2_H_5_) has an EC_50_ of 1.71 nM, while the EC_50_ of the nitro-absent counterpart etodesnitazne (R_1_ = H, R_2_ = OC_2_H_5_) is nearly two orders of magnitude higher (164 nM). Nitazenes containing a halogen, clonitazene (R_1_ = NO_2_, R_2_ = Cl) and flunitazene (R_1_ =NO_2_, R_2_ = F), showed an even higher EC_50_ of 338 and 827 nM, respectively. ^7^ Consistent with these studies, ^7,8^ de Luca et al. showed^11^ that isoto- and metonitazenes have lower K_*i*_ and EC_50_ but similar E_max_ for stimulation of [^35^S]GTP*γ*S binding in rat cortical membranes, compared to fentanyl and DAMGO.

To date, no experimental structure model of *µ*OR in complex with a nitazene is available, hindering the molecular understanding of the pharmacology and structure-activity relationship of nitazenes. Meto-, eto-, proto-, buto-, and isotonitazenes are Schedule I substances under the U.S. Controlled Substance Act. These nitazenes share nitro group as R_1_ and ethyl group as R_3_/R_3’_, while the length of the alkoxy group at R_2_ varies (Figure 1). In this work, we study these nitazenes and three nitro-less analogs (R_1_=H), metodes- and etodesnitazene, which are also Schedule I substances, as well as the commercially available protodesnitazene, which has not been scheduled yet (Figure 1). All eight nitazenes were found to be at least as potent as fentanyl.^7,11^ Accompanying the aforementioned functional study, ^11^ de Luca et al. reported the best scoring poses of isoto- and metonitazenes using Glide^12^ (part of the Schrödinger software suite) and the X-ray cocrystal structure of *µ*OR:BU72 complex (PDB: 5C1M)^13,14^ as a template.

Based on the X-ray structures and cryogenic electron microscopy (cryo-EM) models of *µ*OR in complex with agonist ligands, particularly BU72, fentanyl, and mitragynine pseudoindoxyl (MP), we analyzed the possible binding poses of the aforementioned eight nitazenes (Figure 1). We used consensus docking followed by pose re-ranking with the binding pose metadynamics (BPMD) protocol,^15,16^ as well as refinement with the conventional molecular dynamics (cMD). Two possible binding modes for the nitro-containing nitazenes were found, which differ in positioning of the nitro group, akin to the phenylethyl group of fentanyl in subpocket 1 (SP1) or the indole group of MP in subpocket 2 (SP2). For etonitazene, the SP2-binding mode was estimated to be favored by 2–4 kcal/mol based on the well-tempered metadynamics (WT-metadynamics) simulations^17^ of the refined SP1 and SP2-binding poses. For the nitro-less nitazenes, the SP1-binding mode was ruled out based on the cMD simulations of etodesnitazene bound *µ*OR. Finally, the analysis of putative binding modes of nitazenes and comparison to the experimental structures of other *µ*OR:agonist complexes led to a proposed model for the molecular recognition of opioids by *µ*OR.

## Results and Discussion

### Analysis of the experimental structure models of *µ*OR-ligand complexes

To understand the ligand binding site, we examined the published structure models of all 14 agonist-bound *µ*OR complexes, including one X-ray structure and 14 cryo-EM structure models (Supplemental Table S1). These structures show that the ligand occupies the same central cavity of *µ*OR, and a part of the ligand, e.g., the phenyl group of BU72 or the phenethyl group of fentanyl, reaches into the space between TM2 and TM3, referred to as subpocket 1 (SP1) in Ref,^18^ comprising of Thr120 (Thr^2.57^ in Ballesteros-Weinstein numbering^19^), Val143^3.28^, Ile144^3.29^, and Asp147^3.32^ (Figure 2). The only exception is shown by the cryo-EM model of the mitragynine pseudoindoxyl (MP)-bound *µ*OR (PDB: 7T2G),^18^ in which the methoxyphenyl group extends into the space between TM2 and TM7 (subpocket 2 or SP2,^18^ Figure 2 and S1) comprised of Tyr75^1.39^, Trp318^7.34^, His319^7.35^, and Ile322^7.38^. Considering that the structure of *µ*OR in complex with BU72 (PDB: 5C1M)^13,14^ has the highest resolution (2.1 Å) and the BU72 interaction with SP1 is shared with

**Figure 2:**
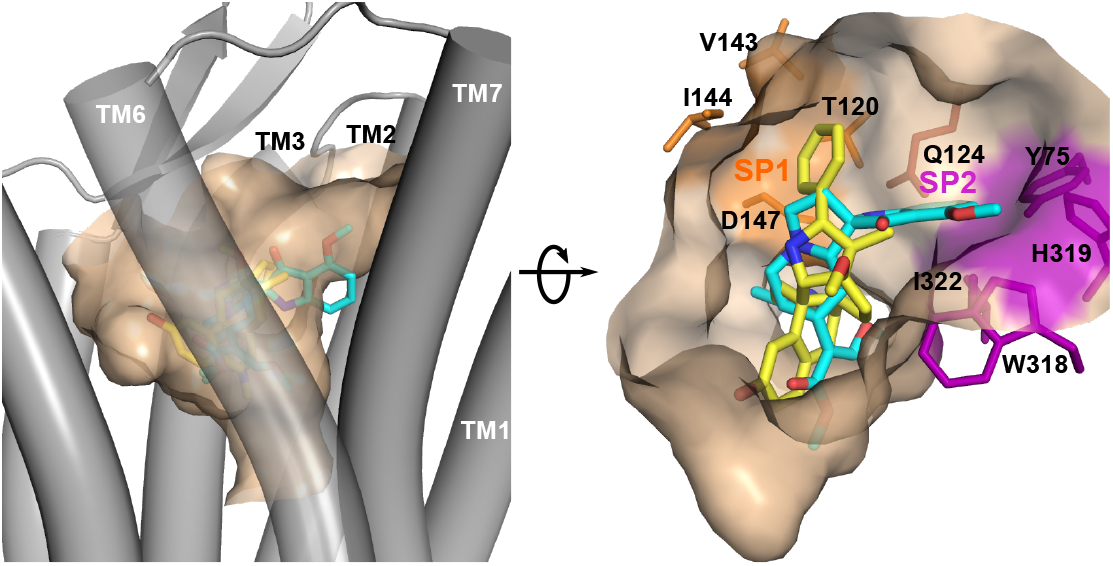
Visualization of the binding pocket of *µ*OR in complex with BU72 (yellow; PDB: 5C1M^13^) or MP (cyan; PDB: 7T2G^18^). For comparison, the BU72 and MP structures are superimposed; but the MP-bound *µ*OR structure is hidden. A main difference is that BU72 reaches into subpocket 1 (SP1, orange) between TM2 and TM3, while MP extends into subpocket 2 (SP2, purple) between TM2 and TM7. SP1 and SP2 are separated by Gln124.

### Validation of the docking and reranking protocol using fentanyl Three binding modes were generated by the docking software

Three docking software, AutoDock Vina, ^21^ Glide^12^ (part of the Schrödinger software suite), and MOE^22^ were utilized to examine whether the binding pose of fentanyl in the recent cryo-EM model (PDB: 8EF5)^20^ can be recapitulated using the BU72-bound *µ*OR structure (PDB: 5C1M)^13,14^ as a template. Binding poses were generated using the three software. For each software, the top three ranked poses were similar in their affinity score (with difference in the second digit or within 1-5% difference) and yet the lig- and conformation was significantly different in some cases (Figure 3). Specifically, the generated poses can be grouped into three categories based on the placement of the propanamide, phenyl, and phenylethyl motifs. The pose is referred to as Match if the positions of all three motifs are similar as in the cryo-EM structure model (PDB 8EF5, Figure 3). The next category is largely similar, except that the locations of the propanamide and phenyl groups are swapped; since phenylethyl is still placed in a hydrophobic region towards the extracellular entrance, this category is dubbed “Akin” (Figure 3). Finally, a third category swaps the positions of the phenylethyl and phenyl groups, resulting in a 180° flip, dubbed “Flipped” (Figure 3). Note, this pose was reported in several previous computational work ^23–25^ and was shown to have similar stability in metadynamics simulations.^26^ It is worth noting that all three software generated at least one Match pose; however, the Glide scoring function cannot distinguish Match from Akin, while the MOE scoring function strongly favors Flipped over Match (Table S2).

**Figure 3:**
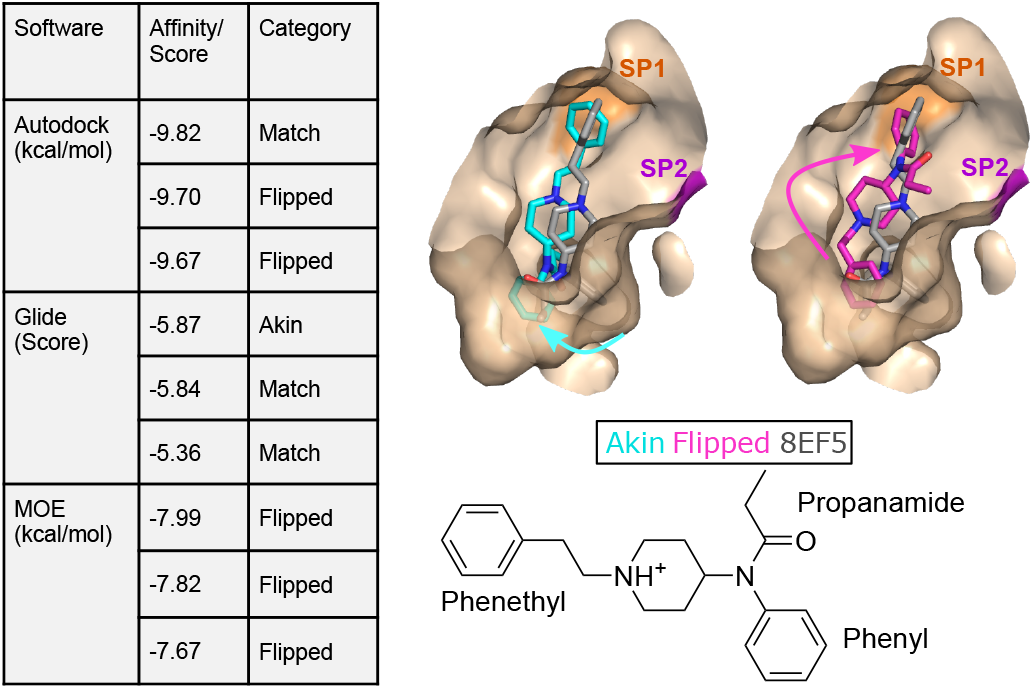
Molecular docking generates three unique poses of fentanyl in *µ*OR. Left: The top three ranked binding poses and their scores given by three docking software. Each pose is labeled as Match, Akin, or Flipped based on the fentanyl position relative to the cryo-EM structure (PDB: 8EF5).^20^ The best score for each pose category is given in Table S2. **Right:** Overlay of the cryo-EM (wheat) and the two docked poses that differ: in Akin (cyan), the phenylethyl group is placed in SP1 between TM2 and TM3, but the propanamide and phenyl groups are swapped in positions; in Flipped (magenta), the phenylethyl and phenyl groups are swapped in positions. For reference the two-dimensional structure of fentanyl is given below.

### The Match pose of fentanyl is most stable in BPMD

It is common knowledge that while the true binding pose is often among the poses generated by a docking program, it is often not the most favorable one based on the docking score.^15^ Since the docking scores for the poses are highly similar, we tested an empirical protocol called binding pose metadynamics (BPMD)^15,16^ to re-rank them by assessing the relative stabilities of the poses in the receptor using metadynamics simulations. Briefly, each docked pose was used to initiate ten independent replicas of 10-ns metadynamics simulations, whereby the ligand RMSD with respect to the docked pose is used as the collective variable (CV), along which a time-dependent biasing potential is deposited to accelerate sampling of ligand conformations with large RMSDs. A fast increase in RMSD would indicate that the initial docked pose is unstable. Furthermore, we monitored the time course of the distance between fentanyl’s piperidine nitrogen and the nearest carboxylate oxygen of Asp147^3.32^, which represents the stability of the conserved salt-bridge interaction. ^13^ The most stable pose should have the lowest average RMSD (over the ten trajectories) and consistently maintain the fentanyl-Asp147^3.32^ salt bridge. Representative poses were tested by first clustering generated poses (separated by software), then the centroid of each cluster was tested. Figure 4 shows the evolution of each quantity for these tested poses, averaged over the ten simulations, and colored by the category the tested pose belonged to. Encouragingly, for the poses generated by all three docking tools, the Match pose consistently had the lowest average RMSD and maintained an average salt-bridge distance of ~3 Å. In contrast, the Flipped or Akin poses consistently had higher average RMSD with some disruption in the conserved salt bridge. Moreover, poses that flip the ligand and place the phenylethyl closer to the center of the receptor pocket had a final average RMSD of 2-3 times larger than the most stable poses. These results suggest that BPMD is capable of differentiating between similarly scored poses, and in this case, finding the experimentally determined pose.

**Figure 4:**
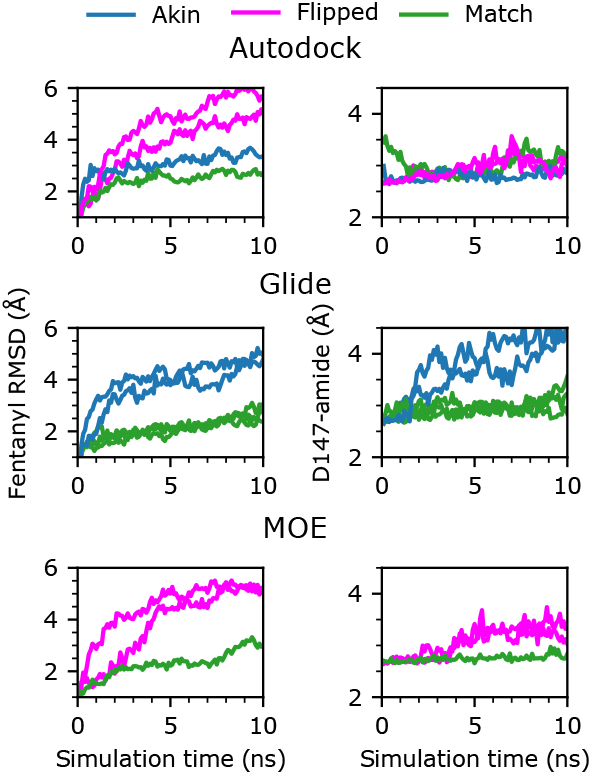
Binding pose metadynamics protocol showed that the Match pose of fentanyl is most stable. Left panel: Time series of the heavy-atom RMSD of fentanyl with respect to the docked pose. **Right panel:** Time series of the minimum distance between the piperidine amine nitrogen of fentanyl and the carboxylate oxygens of Asp147^3.32^. Both RMSD and distance was averaged per frame over ten 10-ns trajectories.

### Determination of the putative binding poses of nitazenes

#### Docking based on the *µ*OR:BU72 cocrystal structure places the nitro group in SP1 or SP3

Having validated the docking and ranking protocol, we proceeded to predict the nitazene binding poses. Eight nitazenes were studied, five of which contain a nitro group at R1 while the remaining three do not (Figure 1). With the constraint of triethylamine and the formation of the Asp147^3.32^ salt bridge, the docked poses from the three software were categorized according to whether the nitro group or the benzene of the benzimidazole motif for the nitro-less nitazenes was placed in SP1 between TM2 and TM3 or another sub-pocket between TM5 and TM6, which will be referred to as SP3 hereafter (Figure 5 and Table S3).

**Figure 5:**
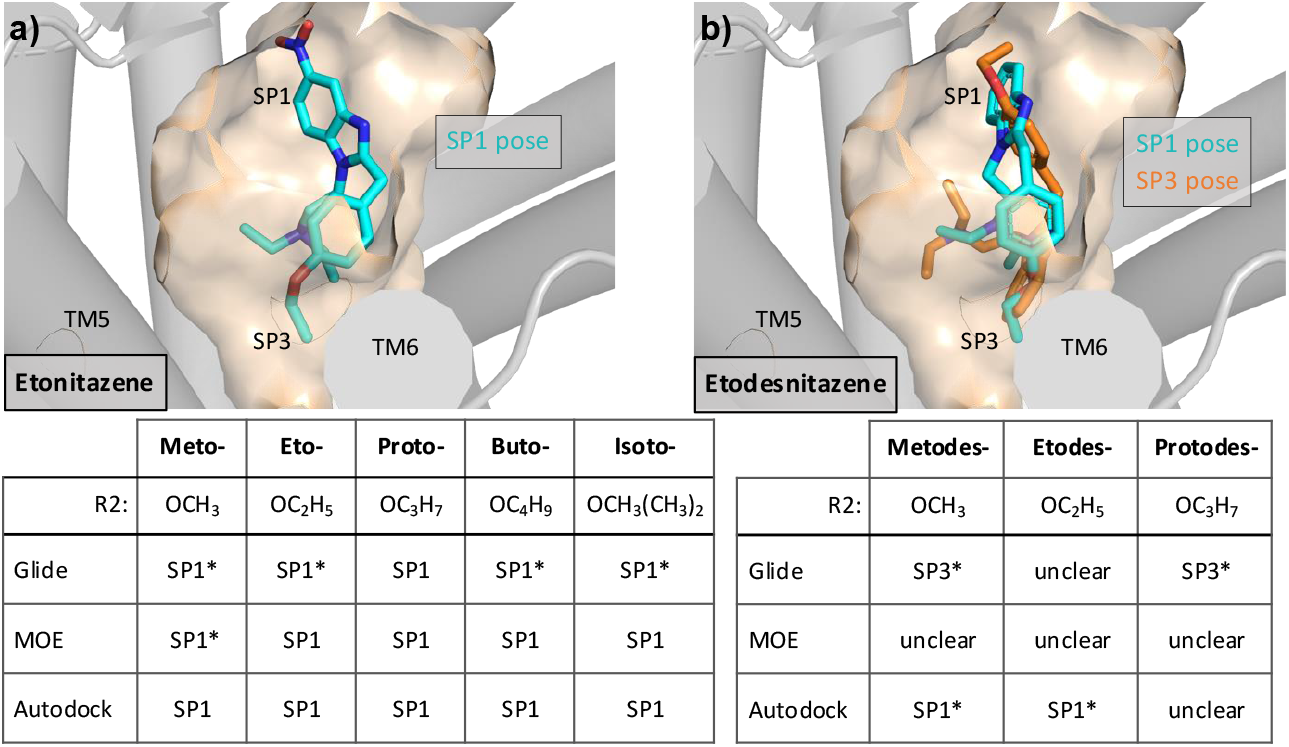
Predicted stable poses of eight nitazenes based on the BU72-*µ*OR co-crystal structure and BPMD. The pose category that was most stable in the BPMD trajectories is given for the nitro-containing (a) and nitro-less nitazenes (b). An asterisk indicates that the software only predicted poses in that particular category.

In the SP1-binding mode, the alkoxy group of the nitazene is placed into SP3, whereas in the SP3-binding mode, it is placed in SP1. In other words, the positions of alkoxy and nitro (or benzene for nitro-less nitazenes) groups are swapped. It is noteworthy that some software only generated the SP1-binding mode for some nitro-containing nitazenes, e.g., Glide generated only the SP1-binding mode for meto-, eto-, buto-, and isotonitazenes (Figure 5a). Curiously, Autodock predicted a third category of poses, dubbed SP2’, where the nitro group was placed on top of Gln124^2.60^ and pointed towards TM1, the *N*-diethyl occupied SP1, and the oxyl tail oriented downward towards the center of the receptor (Figure S4). Thus, for simplicity this pose is not visualized in Figure 5.

#### BPMD suggests the SP1-binding mode as more stable than the SP3-binding mode for nitro-containing nitazenes

To assess which pose, i.e., SP1 or SP3, is more stable, we performed BPMD on the poses generated by the three software for all eight nitazenes. For the nitro-containing nitazenes (meto-, eto-, proto- and butonitazene), the SP1 poses are consistently more stable than the SP3 poses (Figure 5a). For the nitro-less analogs, metodes-, etodes-, and protodesnitazene, however, there is no consistent trend when considering any one compound across the three software or any one software across the three compounds (Figure 5b). For example, considering metodesnitazene, the SP3 pose is favored by two software Glide and Autodock, whereas the SP1 pose is favored by MOE. However, for etodesnitazene, the SP1 pose is favored by two software Glide and Autodock, whereas the SP3 pose is favored by MOE. When considering a single software, e.g., Glide, it only produced SP3 poses for meto- and protodesnitazenes; however, it favored the SP1 pose for etodesnitazene.

#### Docking based on the *µ*OR:MP cryo-EM model places the nitro group in SP2

As our analysis of the experimental structure models of agonist-bound *µ*OR showed that the SP2 between TM2 and TM7 can also be occupied as in the cryo-EM model of the *µ*OR:MP complex (PDB: 7T2G),^18^ we used the latter model to dock the eight nitazenes. Note, the SP1 occupied by BU72, fentanyl, and 11 other ligands and the SP2 occupied by MP are separated by Gln124^2.60^; which can block one of the subpockets due to the movement of the methylamide group (related to a *χ*_2_ angle change). In the BU72-, fentanyl- and 11 other agonist-bound *µ*OR structures (PDB 5C1M and 8EF5, Table S1), the methylamide group of Gln124^2.60^ points down, opening the entrance to SP1 but blocking that to SP2. In contrast, the *µ*OR-MP complex structure 7T2G shows that the methylamide group points up and towards TM3, blocking the entrance to SP1 but opening that to SP2.

Docking of the eight nitazenes based on the cryo-EM model of the *µ*OR:MP complex (PDB 7T2G) resulted again in two distinct binding modes – each determined by whether the nitro group (or the benzimidazole ring) is placed in the opened SP2 or the lower SP3 (Figure 6 and Table S4). In the SP2 mode, the alkoxy group extends towards SP3, whereas in the SP3 mode it points towards SP1. Note, for the nitro-containing nitazenes, the SP3-binding mode was only produced by MOE (Table S4) and its docking score relative to the SP1-binding mode is inconsistent among meto-, eto-, proto-, buto-, and isotonitazenes. Thus, the SP3 binding pose will not be further discussed for the nitro-containing nitazenes. For the nitro-less nitazenes, Glide did not produce the SP3 mode for metodesnitazene and favors the SP2 mode for etodesnitazene. MOE appears to favor the SP2 mode for meto- and etodesnitazenes, while the trend of Autodock is less clear (Table S4). Together, these data suggest that the SP2 mode is slightly favored for meto- and etodesnitazenes.

**Figure 6:**
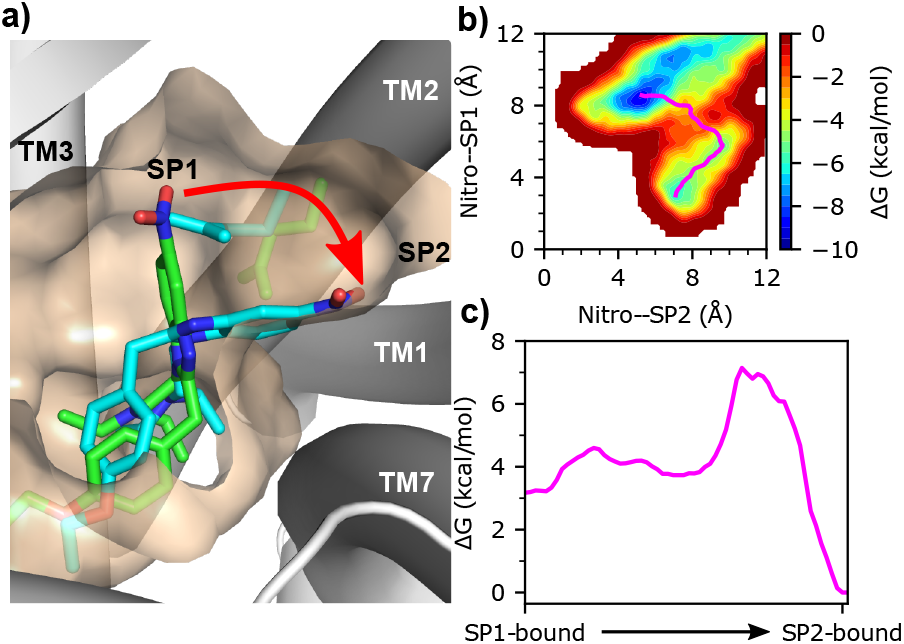
Etonitazene favors the SP2-over the SP1-binding mode. **a)** Visualization of the initial SP1-binding mode (green) and the target SP2-binding mode (cyan) in the metadynamics simulations. **b)** Estimated free energy landscape as a function of the two CVs from a representative metadynamics trajectory. The resulting minimum free energy path from the nudged-elastic band (NEB) calculation is shown in magenta. **c)** Estimated free energy change going from the SP1- to the SP2-binding mode along the minimum free energy path shown in b).

#### Etonitazene favors the SP2-over the SP1-binding mode based on the metadynamics simulations

The SP1 and SP2 poses are mutually exclusive, as neither structure template produced both poses. This means that BPMD cannot be used to evaluate the relative stability between the SP1- and SP2-binding modes. To determine which of two poses (SP1 or SP2) is more favorable, we used etonitazene as a test case and estimated the free energy difference between the two poses using WT metadynamics simulations,^17^ in which a time-dependent biasing potential was deposited on two CVs representing the distance of the nitro group to SP1 or SP2. The CV1 or CV2 is defined as the distance from the nitro nitrogen to the center of mass (COM) of TM2/TM3 or TM2/TM7, respectively (see Methods). A CV1 of 4 Å is representative of the SP1-binding mode, whereas a CV2 of 4 Å is representative of the SP2-binding pose.

Five independent 200 ns metadynamics simulations were performed starting from the refined SP1-binding mode of etonitazene through 100 ns cMD, from which the estimated free energy surfaces as a function of the two CVs were obtained (Figure 6 right). While etonitazene in all five trajectories transitioned out of SP1 within 200 ns, the SP2-binding mode was sampled in three trajectories and was more stable than the SP1-binding mode (Figure S5). In each trajectory, once the nitro group moved away from SP1, the *χ*_2_ of Gln124^2.60^ changed from *trans* to *gauch+* (Figure S6), suggesting that an induced fit mechanism may be responsible. That is, the precise binding mode, SP1 vs. SP2, depends on the ligand structure – etonitazene but not fentanyl induces the movement of Gln124. To understand the transition pathway and estimate the relative stabilities of the SP1/SP2 binding modes, we used the nudged elastic band (NEB) method^27^ to find the minimum energy path in the free energy landscape estimated by metadynamics (Figure 6b and c). The three trajectories that sampled the SP2 pose show that the relative stability of the SP2 vs. SP1 pose is roughly 2-4 kcal/mol (Figure 6c and Figure S5). Note, our focus here is to qualitatively probe the relative stability; a more thorough study is deferred to future work.

#### The SP2-binding mode may be favored by etodesnitazene based on the cMD simulations

We now turn to the nitro-less nitazenes. Unlike the nitro-containing nitazenes, no consensus binding mode was found based on the cocrystal structure *µ*OR:BU72 (PDB: 7C1M) using the three docking programs, e.g., Glide only generated the SP3 mode for metodes- and protodesnitazene, while Autodock only generated the SP1-binding mode for metodes- and etodesnitazene (Figure 2 and Table S3). The BPMD simulations of the SP1 and SP3 binding poses for cases where both poses were predicted by the docking software were also inconclusive (Figure 2 and Figure S3). However, when comparing the SP2 and SP3 poses obtained from the *µ*OR:MP structure based docking, the SP2 pose appears to be somewhat preferred for meto- and etodesnitazenes (Table S4).

To test which binding pose is most favorable for nitro-less nitazenes, we used etodesnitazene as a model and conducted cMD simulations starting from the top docked poses of the SP1, SP2, and SP3 binding modes. Three independent 250-ns simulations were conducted for each pose. To assess the stability of each pose, we examined the RMSD with respect to the docked pose (Figure 7). Starting from the SP1 and SP3 docked poses, the RMSD rapidly increased to 3–6 Å in all three replica runs, whereas the RMSD in two out of three runs starting from the SP2 pose remained below 2 Å(Figure 7). It is noteworthy that in one trajectory starting from the SP2 pose, the RMSD increased to 4.5 Å and stayed for approximately 75 ns before decreasing below 2.5 Å and remaining so in the last 150 ns (Figure 7).

**Figure 7:**
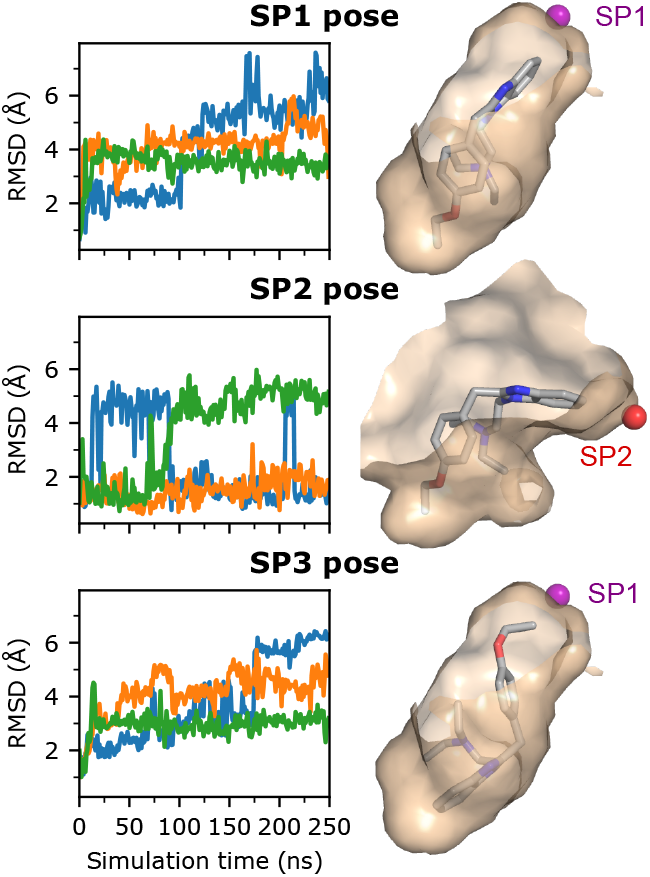
The SP2-binding appears to be more stable than the SP1- and SP3-binding modes for etodesnitazene. Left panel. Time evolution of the heavy-atom RMSD of etodesnitazene with respect to the SP1 (top), SP2 (middle) or SP3 (bottom) docked pose. For each docked pose, three independent 250-ns simulations (blue, orange, and green) were conducted. **Right panel**. Visualization of the SP1-(top), SP2-(middle), and SP3 (bottom) docked poses. The reference points for SP1 and SP2 are shown as purple and red spheres, respectively, and defined as the center of mass of the subpocket (see Methods).

To further compare the relative stabilities of the three poses, we examined the time series of the ligand-pocket distance. For the SP1 and SP2 pose, this is defined as the distance between the carbon adjacent to the nitro group to the COM of the subpocket (see Methods), while for the SP3 pose, the distance between the terminal carbon of the ethoxy tail and the COM of SP1 is used. In simulations initialized from the SP1 docked pose, benzimidazole consistently moved further away from SP1 (Figure S7). Curiously, the *χ*_2_ angle of Gln124^2.60^ changed from *trans* to primarily *gauche+* after 20 ns in all three simulations (Figure S7), similar to that of etonitazene as it transitioned from the SP1 to the SP2-binding mode (Figure 6 and Figure S6). This change in the *χ*_2_ angle allows the carboxamide group to move into SP1, pushing the benzimidazole group out of SP1. Thus, it appears that in the absence of a nitro group the benzimidazole is unable to keep Gln124^2.60^ out of SP1. Similarly, in simulations starting from the SP3 pose, the ethoxy tail moves out of SP1 in two of the three simulations (Figure S7), while in the one simulation where the tail remained in SP1, Gln124^2.60^ freely transitions between *trans, gauch+*, and *gauch-* (Figure S7). By contrast, in two of the three simulations initialized from the SP2 pose, etodesnitazene stayed or only briefly moved out before returning to the SP2 pose (Figure S7), consistent with the time evolution of the ligand RMSD (Figure 7). These data suggest that the SP2-binding mode is more stable than the SP1- and SP3-modes; however, further investigation is needed to confirm.

### A putative model of *µ*OR:nitazene recognition and comparison to fentanyl and MP

#### Ligand can access the hydrophobic SP1 or the hydrophilic SP2 mediated by the movement of Gln124^2.60^

After establishing the SP2-binding mode as the most favorable one for etonitazene, we proceeded to refine the docked SP2 poses of meto-, eto-, proto-, buto-, and isotonitazenes using 100 ns cMD simulations. These compounds were tested, as they are the most potent nitazenes.^7,8^ Note, the simulation of etonitazene was continued to 300 ns and no visible position or conformational change of the ligand was observed (Figure S8). We then compared the binding poses of these nitazenes with those of fentanyl and MP from the cryo-EM structures in an attempt to understand the general trend in molecular recognition between *µ*OR and opioids with different chemical scaffolds. Note, the nitro-less nitazenes are excluded from the following discussion, as their binding modes require further investigation in the future.

Figure 8a visualizes how fentanyl, MP, and etonitazene bind within the receptor pocket.

**Figure 8:**
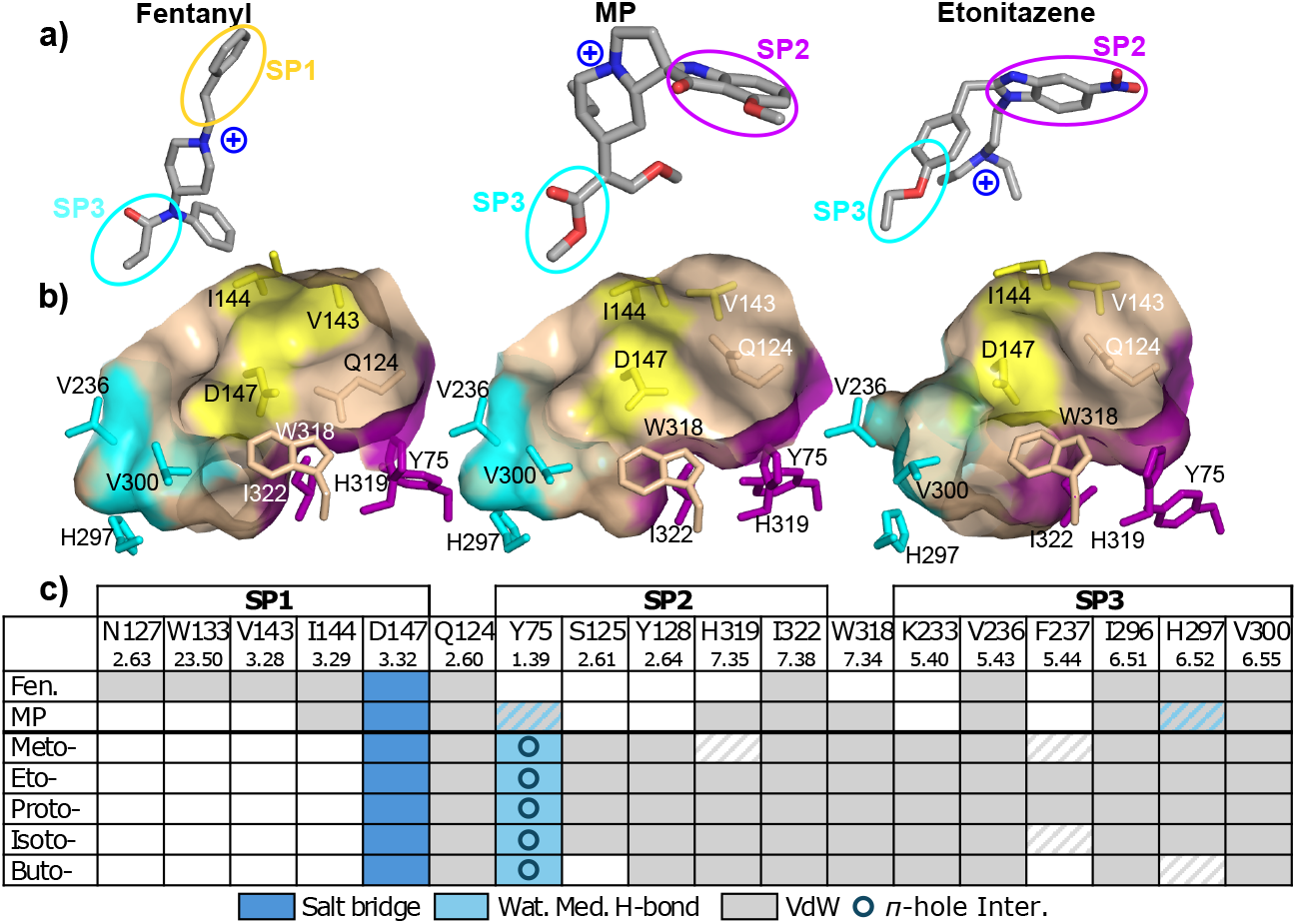
Comparison of the receptor binding modes of nitro-containing nitazenes with fentanyl and MP. **a)** Three-dimensional structures of fentanyl (PDB 8EF5), ^20^ MP (PDB 7T2G),^18^ and etonitazene. **b)** A surface view of SP1 (yellow), SP2 (purple), and SP3 (cyan) residues interacting with fentanyl (left), MP (middle) and etonitazene (right). **c)** Contact analysis of fentanyl, MP, and nitro-containing nitazenes using the final 50 ns simulations. Van der Waals contacts (gray) and salt-bridge interactions (dark blue) are defined using a heavy atom distance cutoff of 4.5 Å and with an occupancy above 50% (for the nitazenes). Gray strips indicate van der Waals contacts with occupancies of 20–50%. Water-mediated hydrogen bonds (light blue) with occupancies above 20% are included. Circle indicates the *π*-hole interaction. Blue strips indicate possible water-mediated hydrogen bonds based on the cryo-EM model (PDB 7T2G).^18^ Detailed maps of receptor-ligand interactions for the nitro-containing nitazenes are given in Figure S10 and S11. A complete list of receptor-ligand contacts with occupancies is given in Table S6.

All three utilize a tertiary amine as part of their scaffolds to form a salt bridge with Asp147^3.32^, which anchors the molecule in the central cavity of the receptor. In the cryo-EM model of *µ*OR:fentanyl complex (PDB 8EF5), ^20^ fentanyl is oriented vertically, whereby the phenethyl group occupies the hydrophobic SP1, forming hydrophobic contacts with Val143^3.28^ and Ile144^3.29^ between TM2 and TM3. In contrast, the hydroxyl-substituted indole of MP and the nitro-substituted benzimidazole of the nitro-containing nitazenes occupy the more hydrophilic SP2, interacting with Tyr75^1.39^, His319^7.35^, and Ile322^7.38^. Note, Gln124^2.60^ separates SP1 and SP2 and blocks a portion of the unoccupied subpocket; for fentanyl it rests on the bottom of SP2 making the region more shallow, while for MP and the nitro-containing nitazenes it orients towards Val143^3.28^ blocking access to SP1.

#### The nitro group forms the water-bridged hydrogen bonds and *π*-hole interaction with SP2

The cMD refinement simulations of the nitro-containing nitazenes in the SP2 pose showed that the nitro group forms a water-mediated hydrogen bond with Tyr75^1.39^, whereby either oxygen of the nitro group acts as a hydrogen bond acceptor for a water molecule while the hydroxyl oxygen of Tyr75^1.39^ acts as a donor. With lower (less than 20%) occupancies, water-mediated hydrogen bonds were also formed with Gln124^2.60^, Trp318^7.34^, and His319^7.35^. Regarding the stabilization of MP in SP2, the cryo-EM model (PDB 7T2G^18^) suggests that the methoxy oxygen of MP may form a water-mediated hydrogen bond with the hydroxyl oxygen of Tyr75^1.39^, similar to the nitro group of the tested nitazenes (Figure 9a).

**Figure 9:**
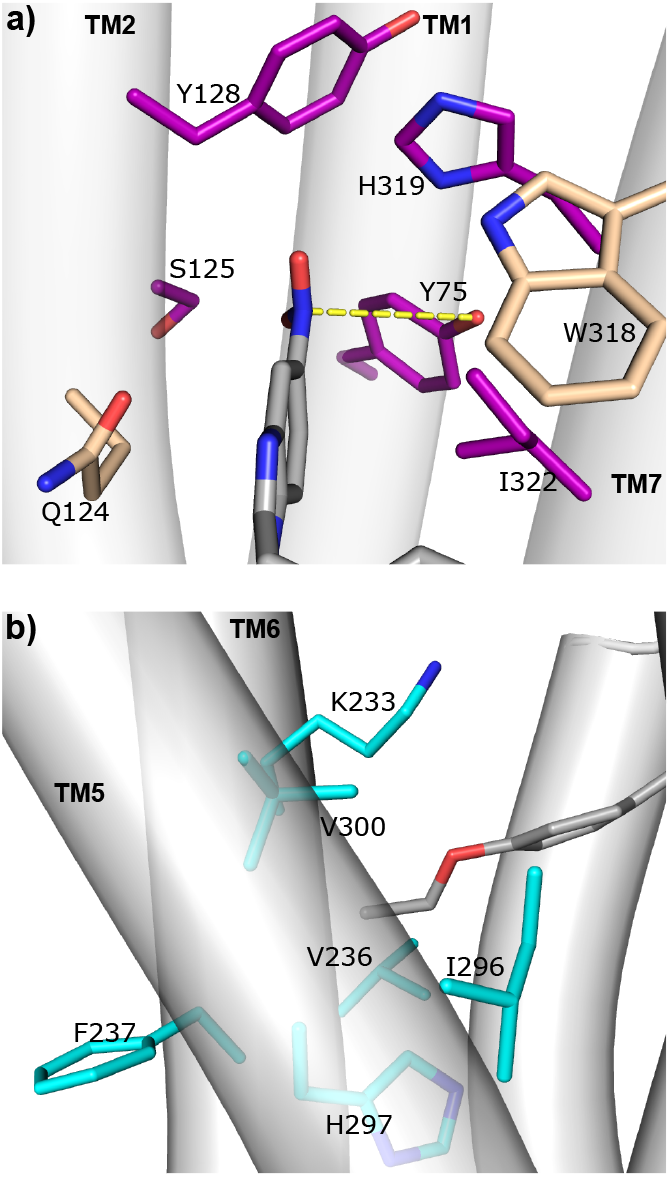
Etonitazene interactions with the residues in SP2 and SP3. A zoomed-in view of a simulation snapshot showing the interactions of etonitazene with the residues in SP2 (**a**) and SP3 (**b**). TMs are represented by cylinders and labeled. The *π*-hole interaction between the nitro group and Tyr75^1.39^ is high-lighted in yellow.

Interestingly, the nitro group’s nitrogen in the tested nitazenes is consistently within a van der Waals distance (below 4.5 Å) from the hydroxyl oxygen of Tyr75^1.39^. For example, in the last 100-ns simulation of the *µ*OR:etonitazene complex, the average N–O distance was 3.8 Å while this distance was below 4.5 Å 88% of the time. This is intriguing, as the nitro group’s electron-deficient nitrogen is known to form a stabilizing *π*-hole interaction with the lone pair of oxygen,^28^ although such an interaction is not explicitly described by the molecular mechanics force field of nitrobenzene^29^ used in this work. We hypothesize that this *π*-hole interaction helps stabilize the nitro group in SP2 (Figure 9a). For etodesnitazene and other nitro-less nitazenes, neither *π*-hole nor water-mediated interaction is possible, which may explain the significantly lower binding affinity and potency of the - des analog compared to the parent nitro-containing nitazene.^8^

#### The alkoxy tail forms hydrophobic contacts with SP3 residues

Similar to fentanyl and MP, the nitazenes make van der Waals contacts with the residues flanking SP3, including Val236^5.43^ on TM5 and Ile296^6.51^, His297^6.52^, and Val300^6.55^ on TM6 (Figure 8c and 9b). Interestingly, all five nitazenes form hydrophobic contacts with part of the sidechain of Lys233^5.40^ on TM5, while all but butonitazene interact with the backbone of Phe237^5.43^ on TM5; in contrast, these interactions are absent with fentanyl and MP (Figure 8c and 9b). Butonitazene has the longest alkoxy tail among the five nitro-containing nitazenes. During the cMD refinement simulation, the butoxy tail extended upward along TM5, gaining hydrophobic contacts with Glu229^5.36^ and Leu232^5.39^, which lie above SP3 (Figure S9). However, regardless of this configuration, butonitazene still interacts with the residues flanking SP3, namely Val236^5.43^, Ile296^6.51^, and Val300^6.55^(Figure 8c and Figure S9). This upward movement disrupts butonitazene’s interaction with the backbone of Phe237^5.43^ and significantly weakens its interaction with His297^6.52^, which are residues at the base of SP3 (Figure 8c and Figure S9). Given the very limited sampling in the refinement cMD simulations, our analysis of the SP3 interactions of butonitazene as well as meto-, proto-, and isotonitazenes should be taken with a grain of salt. A more thorough study will be conducted in the future.

We wondered if the alkoxy group might be stabilized by water-mediated interactions with His297^6.52^. In the X-ray structure of the BU72-bound *µ*OR (PDB 5C1M),^13^ two water bridge the interaction between N*δ* of His297^6.52^ and the hydroxyl oxygen of BU72. Our previous meta-dynamics simulations suggested that His297^6.52^ may directly interact with fentanyl and analogs in the ligand-*µ*OR unbinding process.^26,30^ Although water is not resolved in the cryo-EM structure of MP-bound *µ*OR (PDB 7T2G),^18^ the distance between the methoxy oxygen and either of the His297^6.52^ imidazole nitrogens is less than 5 Å, suggesting that water-bridged interactions may be possible. In the cMD refinement simulations of the five nitazenes, water-bridged interactions between the alkoxy group and His297^6.52^ were not observed, although conceivable based on the distance.

## Conclusions

In this work, we determined the putative nitazene-*µ*OR interactions using docking and MD tools. We first tested a protocol comprised of consensus docking based on the *µ*OR:BU72 X-ray structure (PDB 5C1M)^13,14^ followed by BPMD. This protocol was able to recapitulate fentanyl’s binding pose in the cryo-EM model (PDB 8EF5),^20^ in which phenethyl occupies a subpocket SP1. We next applied the protocol to determine the binding poses of five nitro-containing and three nitro-less nitazenes based on the aforementioned *µ*OR:BU72 X-ray structure (PDB 5C1M)^13,14^ as well as the *µ*OR:MP cryo-EM model (PDB 7T2G),^18^ in which MP occupies an adjacent subpocket SP2 while SP1 (occupied by fentanyl in PDB 8EF5) is blocked by Gln124^2.60^.

Docking based on the *µ*OR:BU72 template generated the SP1 and SP3 binding modes, while the SP2-binding mode was generated by the *µ*OR:MP template. Subsequent BPMD and metadynamics simulations suggest that the SP2-binding mode is most stable for etonitazene and likely also meto-, proto-, isoto-, and butonitazenes. The cMD data suggest that the SP2-binding mode is stabilized by the water-mediated hydrogen bond interactions between the R1 nitro group of the nitazenes and Tyr75^1.39^ of the receptor. Our results disagree with those of de Luca et al.,^11^ which suggest the SP3 binding mode for isoto- and metonitazenes, whereby the nitro group interacts with Lys303 on TM5. The preference for the SP3 binding mode may be related to the use of the *µ*OR:BU72 structure template and Glide’s flexible docking feature. Regarding the nitro-less nitazenes, the cMD simulations of etodesnitazene showed that, unlike fentanyl’s phenethyl group that occupies SP1, the unsubstituted benzimidazole group is unable to displace Gln124^2.60^ from SP1, making the SP1-binding mode unfavorable. The cMD data also provided evidence to support the SP2-binding mode; however, further investigation is needed to confirm.

Our computational analysis of the putative binding modes of the nitro-containing nitazenes and comparison to the X-ray or cryo-EM binding modes of BU72, fentanyl, and MP led us to propose a putative model for *µ*OR-opioid recognition. In this model, the ligand is anchored to TM3 through a salt bridge between the charged tertiary amine located near the center of the structure and Asp147^3.32^, while the substituents R1, R2, and R3 (Figure 1) are recognized by the subpockets: SP1 (between TM2 and TM3), SP2 (between TM2 and TM7), and SP3 (between TM5 and TM6). Separated by Gln124^2.60^, the hydrophobic SP1 and hydrophilic SP2 are alternately accessible; the former prefers an aromatic moiety, e.g., the phenyl ring in BU72 or fentanyl, and the latter favors a polar group, e.g., the methoxy group in MP or the nitro group of nitazenes, which can form a water-mediated hydrogen bond interaction with Tyr75^1.39^. The SP3 is largely hydrophobic; however, it contains His297^6.52^, which may stabilize a polar group via water-mediated hydrogen bond interactions, e.g., with the methoxy tail of MP or the alkoxy tail of the nitazenes.

The current work has several caveats that are worth mentioning. To corroborate the conclusion regarding the relative stability of the SP2 vs. SP1 pose, additional metadynamics simulations initiated from alterative poses may be desirable. However, a potential issue is the gateway behavior of Gln124^2.60^, which occupies SP1 when the pocket is not occupied by a ligand. During our metadynamics simulations, Gln124^2.60^ transitions from *trans* to *gauche+* when etonitazene moves out of the SP1 region and does not transition back to *trans* for the remainder of the simulation (Figure S6). Therefore, starting from an alternative pose (e.g., SP2 pose) may require biasing the conformation of Gln124 in order to sample the SP1 pose, which is beyond the scope of the current work. For the nitro-less nitazenes, while the SP1-binding mode is ruled out and the SP2-binding mode appears most stable for etodesnitazene, the stability of the SP3 binding mode, especially for other nitro-less nitazenes, warrants further investigation. Another caveat is the relatively short (100-ns) simulation length for pose refinement, although extending the simulation to 300 ns did not result in a significant shift in the position of etonitazene. We should also note that the simulations were conducted without the G-protein complex, which stabilizes the activated state;^13^ simulations on a longer timescale might risk conformational changes away from the active state. Despite these limitations, our study provides valuable insights into the molecular recognition of nitazenes and other opioids by the *µ*OR, laying the foundation for detailed investigations into the structure-activity relationships.

## Methods and Protocols

### Molecular docking of fentanyl

The molecular docking of fentanyl was conducted based on the co-crystal structure of *µ*OR:BU72 complex (PDB: 5C1M)^13^ after reversing the modification to Cys57. The nanobody mimicking the G-protein complex was removed, as well as any lipids. The three-dimensional structure of fentanyl was downloaded from PubChem; the piperidine nitrogen of fentanyl was protonated, using the tools in Maestro (before using Glide) or MOE or PyMol v2.5 (before using Autodock-Vina).

#### Docking with Glide

*µ*OR was prepared using the protein preparation wizard^31^ in Maestro and fentanyl was prepared using LigPrep. A docking grid centered around BU72 was created, with a distance constraint between the piperidine nitrogen of fentanyl and Asp147^3.32^. This grid consisted of two parts: a 10×10×10 Å inner box that constrained the center of the docked ligand, and an outer box that was used to compute the grid potentials. The size of the outer box was determined automatically using the size of BU72. A maximum of 10 poses was generated using Glide^12^ using standard precision (SP).

#### Docking with MOE

*µ*OR was first prepared within MOE2022.02, ^22^ then docking was initiated after identifying the binding pocket based on the position of BU72 and setting a pharmacophore filter to ensure a hydrogen bond with Asp147. 50 poses were generated using London dG scoring function and the triangle matcher method for placement, with 25 being refined using GBVI/WSA dG function.

#### Docking with AutoDock Vina

Both *µ*OR and fentanyl were prepared using Autodock-Tools.^32^ All nonpolar hydrogens were removed. Docking was confined to a 20×20×25 Å box centered on BU72; exhaustiveness was set to 250, maximum number of poses to 100, and the affinity range to 10 kcal/mol. Generated poses were filtered to ensure a salt bridge with Asp147, defined by a distance cutoff of 4.0 Å between the piperidine amine nitrogen of fentanyl and the nearest carboxylate oxygen of Asp147 and ensuring that the amine hydrogen is oriented towards the carboxylate. Version 1.2.3 of AutoDock Vina ^21,33^ was used.

### Molecular docking of nitazenes

Molecular docking of nitazenes was performed using either the *µ*OR:BU72 complex (PDB: 5C1M^34^) or the *µ*OR:MP complex (PDB: 7T2G^18^). The former was prepared as described for fentanyl, while the latter was prepared in a similar manner (i.e. removal of the G-protein complex and resolved ligands). The three-dimensional structure of each tested nitazene was taken from PubChem; the structure of protodesnitazene was generated by modifying the three-dimensional structure of protonitazene as there was no existing entry for protodesnitazene. The docking procedure and parameters for fentanyl outlined above were used for nitazenes.

### Binding pose metadynamics (BPMD) for pose evaluation

#### Selecting poses

For most compounds Glide generated less than the maximum ten poses. The five best scoring poses (or all poses if less than five were generated by Glide) were used to initiate BPMD simulations. In contrast, MOE and Autodock generated a larger amount of poses. To narrow down the poses for evaluation, each pose set (excluding poses with an unfavorable, positive score) was clustered using hierarchical clustering (as implemented in CPPTRAJ^35^) based on the pairwise root-mean-square deviation (RMSD) of the ligand. Clustering was conducted until clusters were at least 2 Å RMSD different from each other. The centroid of each cluster was then used to initialize BPMD simulations.

#### System preparation

Each selected pose was prepared using CHARMM-GUI.^36^ A pure POPC bilayer composed of approximately 175 lipids per leaflet was created, then *µ*OR with fentanyl or a nitazene derivative in the selected pose was inserted into the bilayer using the PPM 2.0 webserver. ^37^ Water molecules were added with a layer thickness of 22.5 Å; sodium and chloride ions were added to neutralize the system (based on the protonation states determined in Ref.^25^) and reach an ionic strength of 150 mM.

#### Force field parameters

The protein and lipids were represented by CHARMM36m^38^ and CHARMM36 lipid^39^ force fields, respectively. Water was presented by the modified TIP3P model^40,41^ and ions were represented by the CHARMM sodium/chloride force field.^42^ The nitazenes studied in this work were parameterized using CGenFF,^43,44^ which assigns a small molecule with CHARMM parameters by searching for similar molecules or fragments that have been explicitly parameterized based on bond perception and atom typing. In particular, the parameters of the nitro group were based on those of nitrobenzene.^29^ All nitazene compounds demonstrated CGenFF penalty scores below the threshold of 50, indicating that the default parameters may be sufficiently accurate and no additional parameter optimization is required.^44^

#### Settings of MD

Following the recommended protocol in CHARMM-GUI,^36^ the system was minimized over 5000 steps then heated to 300 K over 125 ps with restraints on protein positions, lipid positions, and lipid dihedrals. These restraints were then gradually reduced in steps over 2.25 ns. OpenMM^45^ v8.0 was used for both the recommended protocol and metadynamics simulations. A cutoff of 12 Å was used for nonbonded interactions, while long-range electrostatics were calculated using Particle Mesh Ewald summation with the grid size of 1 Å and real-space cutoff of 10 Å. Pressure of 1 bar was applied using the Monte-Carlo barostat^46^ with a coupling frequency of 100 steps, with the x and y dimensions coupled isotropically. Unless otherwise specified, temperature was set to 300 K using the Langevin thermostat^47^ with a friction coefficient of 1. SETTLE^48^ was used for water molecules while SHAKE^49^ was used for all other atoms.

#### Settings of metadynamics

For BPMD, we followed the protocol of Ludauskis and Gervasio et al.^16^ The metadynamics class in OpenMM was used to perform WT-metadynamics. ^17^ To allow SETTLE^48^ and SHAKE^49^ algorithms were used for constraining bonds and angles involving hydrogen atoms in water and protein, respectively. A hydrogen mass repartition scheme^50^ was applied to the system to allow a timestep of 4 fs. A flat-bottom restraint with a constraint of zero was applied to fix a periodic boundary issue.^16^ Ten 10 ns WT-metadynamics ^17^ simulations were performed for each pose, where the collective variable (CV) was set to the RMSD of the ligand heavy atom positions with respect to those in the first frame of the simulation. The metadynamics parameters were identical to those by Ludauskis et al.:^16^ a bias was deposited every 1 ps with a hill height of 0.3 kcal/mol and a width of 0.02 Å, and the bias factor was set to 4. The bias was applied to a grid spanning 0 to 10 Å, with five points per bias width.

Each pose was evaluated according to the time evolution of the ligand RMSD and the minimum distance between the charged amine nitrogen and the carboxylate oxygen of Asp147^3.32^; these quantities were calculated for each frame of one trajectory followed by averaging over the ten independent trajectories.

### Metadynamics for estimating relative stabilities of the SP1- and SP2-binding modes of etonitazene

To sample transitions from SP1- to SP2-binding modes, we first refined etonitazene in SP1 through cMD according to the protocol detailed above. SP1 was selected as we hypothesized that once the nitro group exits the sub-pocket and moves into the main pocket, Gln124 will move into SP1 thus opening the pathway to SP2. We conducted five 200 ns WT-metadynamics simulations of *µ*OR:etonitazene complex starting from the cMD refined SP1-binding pose. Two CVs were used to represent the progress towards the SP1- and SP2-binding modes. For the former, we used the distance between the nitro group nitrogen and the C*α* center of mass (COM) of residues 122-126 on TM2 and 140-144 on TM3. For the latter, we used the distance between the nitro nitrogen and C*α* COM of residues 122-126 on TM2 and 319-323 on TM7. In WT-metadynamics, biasing deposits were added every 50 ps with a hill height of 0.25 kcal/mol and a width of 0.25 Å, and a bias factor of 14. PLUMED 2.7^51^ was used to run WT-metadynamics in OpenMM and calculate the approximate free energy landscape along the CVs. Pathways from the SP1- to the SP2-binding modes were produced using the nudged elastic band method through a modified script from the R package metadynminer.^52^

### Conventional MD (cMD) for binding pose refinement and analysis

#### Nitro-containing nitazenes

cMD was used to refine the SP2-binding mode for nitro-containing nitazenes. For each molecule, the system was prepared as for initializing BPMD trials followed by the 100-ns (300-ns for etonitazene) cMD simulation using OpenMM^45^ v8.0. The aforementioned parameters and settings in MD were used.

*µ*OR-ligand interaction analysis was performed on the last 50 ns of the 100 ns cMD simulation using MDTraj^53^ with a heavy-atom distance cutoff of 4.5 Å for hydrophobic contacts and salt bridges, while water-mediated interactions were analyzed using MDAnalysis. ^54^ All other analysis, such as RMSD calculations, were performed with CPPTRAJ^35^

#### Etodesnitazene

cMD simulations were performed to test the SP1, SP2, and SP3 binding modes of etodesnitazene. Three independent simulations were conducted for each binding mode starting from the docked pose, each lasting 250 ns. Measured distances were for the analysis of whether a motif was in the subpocket, using either the benzimidazole carbon at the 6-position (which connects to the nitro of nitro-containing nitazenes) or the terminal carbon of the etoxy motif. The reported distance refers to that between this carbon and the COM of C*α* atoms of either residues 122-126 and 140-144 (SP1), or residues 122-126 and 319-323 (SP2). Distances, RMSDs, and dihedrals were calculated using CPPTRAJ^35^ while water-mediated hydrogen bonds were analyzed using MDAnalysis. ^54^ The contacts shown in Figure 8 were found using the final 50 ns of the SP2 pose trajectory indicated in blue in Figure S7; this was done as other nitazene contacts were found using a single trajectory.

## Data Availability

The docked and refined poses of all nitazenes as well as all simulation inputs are freely downloadable from GitHub (https://github.com/janashen/Nitazenes).

### Acknowledgement

Joseph Clayton is supported by the ORISE fellowship, which is a Research Participation Program at the FDA administered through the Oak Ridge Institute for Science and Education (ORISE) under the agreement between the FDA and Department of Energy. J.S. is supported by the National Institutes of Health (R35GM148261 and R01CA256557).

## Disclaimer

This article reflects the views of the authors and should not be construed to represent FDA’s views or policies. The mention of commercial products, their sources, or their use in connection with material reported herein is not to be construed as either an actual or implied endorsement of such products by the FDA.

## Supporting Information Available

Table S1 containes the published X-ray structures and cryo-EM models of *µ*OR in complex with agonist ligands. Table S2–S4 contain the docking scores of fentanyl and nitazenes from the three software programs based on two different templates. Table S5 contains the occupancies of the water-mediated hydrogen bonds between the nitro-containing nitazenes and *µ*OR. Fig. S1 compares the experimentally determined binding poses of BU72, DAMGO, fentanyl, and MP. Fig. S2 and S3 contains the BPMD pose evaluation for the nitro-containing (Fig. S2) and nitro-less (Fig. S3) nitazenes. Fig. S4 details the SP2’ pose predicted only by Autodock. Fig. S5 contains the approximate free energy surfaces for the transition between SP1 and SP2-binding modes of etonitazene from WT-metadynamics simulations. Fig. S6 contains the time evoluation of CVs and Gln124^2.60^ *χ*_2_ angle in the WT-metadynamics simulations of *µ*OR:etonitazene complex. Fig. S7 contains the time evolution of pocket distances, Gln124^2.60^*χ*_2_ angle, and etonitazene RMSD in the etodesnitazene cMD simulations. Fig. S8 contains a visualization of an extended etonitazene cMD simulation and the time evolution of the RMSD with respect to the final frame. Fig. S9 contains the time evolution that suggests the possible interaction between the nitro-containing nitazenes and Tyr75^1.39^.

