## Supporting Information for "A Putative Binding Model of Nitazene Derivatives at the *μ*-Opioid Receptor"

Joseph Clayton,[†,‡] Lei Shi,[¶] Lidyia Stavitskaya,[*,§] and Jana Shen[*,‡]

†*Division of Applied Regulatory Science, Office of Clinical Pharmacology, Office of Translational Sciences, Center for Drug Evaluation and Research, United States Food and Drug Administration, Silver Spring, Maryland 20993, United States*

‡*Department of Pharmaceutical Sciences, University of Maryland School of Pharmacy, Baltimore, MD 21201, United States*

¶ *National Institutes on Drug Abuse 251 Bayview Blvd, Baltimore, MD 21224, United States*

§*Division of Applied Regulatory Science, Office of Clinical Pharmacology, Office of Translational Sciences, Center for Drug Evaluation and Research, United States Food and Drug Administration, Silver Spring, MD 20993, United States*

### Supplemental tables

Table S1: X-ray and cryoEM structure models of agonist-bound $\mu$OR in the PDB sorted by the overall resolution.

| Active $\mu$OR (mouse) | | | | |
|---|---|---|---|---|
| Ligand no. | Entry | Resolution (Å) | Agonist | Method |
| 1 | 5C1M | 2.1 | BU72 | X-ray |
| 2 | 7T2G | 2.5 | MP | cryoEM |
| 3 | 7SBF | 2.9 | PZM21 | cryoEM |
| 4 | 7T2H | 3.2 | Lofentanil | cryoEM |
| 5 | 7U2L | 3.2 | C6-guano | cryoEM |
| | 7U2K | 3.3 | C6-guano | cryoEM |
| 6 | 6DDF | 3.5 | DAMGO | cryoEM |
| Active $\mu$OR (human) | | | | |
| 7 | 8EFO | 2.8 | PZM21 | cryoEM |
| 8 | 8EF6 | 3.2 | Morphine | cryoEM |
| 9 | 8EFB | 3.2 | Oliceridine | cryoEM |
| 10 | 8EFL | 3.2 | SR17018 | cryoEM |
| 11 | 8F7Q | 3.2 | $\beta$-endorphin | cryoEM |
| 12 | 8EF5 | 3.3 | Fentanyl | cryoEM |
| 13 | 8EFQ | 3.3 | DAMGO | cryoEM |
| 14 | 8F7R | 3.3 | Endomorphin | cryoEM |

The BU72- and MP-bound structures (red) were used as templates for our docking campaign. The fentanyl-bound structure (blue) was used for validation of the docking and scoring protocol.

Table S2: Docking poses and scores of fentanyl based on the X-ray structure of BU72-$\mu$OR complex.

| Software | Match | Akin | Flipped |
|---|---|---|---|
| Autodock | -9.82 (kcal/mol) | -9.70 (kcal/mol) | -9.67 (kcal/mol) |
| Glide | -5.84 | -5.87 | — |
| MOE | -5.63 (kcal/mol) | — | -7.99 (kcal/mol) |

Note, only poses that show a potential salt bridge between the piperidine and Asp147 are retained. A hydrogen bond constraint was added during docking in both MOE and Glide; for Autodock a similar constraint was added after docking by imposing a distance cutoff of 4 Å between the piperidine nitrogen and the nearest carboxylate oxygen of Asp147, and ensuring the hydrogen on the piperidine nitrogen was closer to the carboxylate oxygen than the piperidine nitrogen.

Table S3: Docking poses and scores of 8 nitazenes based on the X-ray structure of $\mu$OR:BU72 complex

| | Metonitazene | | Etonitazene | | Protonitazene | | Isotonitazene | |
|---|---|---|---|---|---|---|---|---|
| Software | SP1 | SP3 | SP1 | SP3 | SP1 | SP3 | | |
| Glide | -7.2 | — | -7.4 | — | -4.9 | -5.6 | -6.6 | — |
| MOE | -8.7 | -7.7 | -8.7 | -7.2 | -9.1 | -8.5 | -9.3 | -9.0 |
| Autodock | -7.8 | -7.8 | -8.7 | — | -8.6 | -8.1 | -8.7 | -7.6 |
| | **Butonitazene** | | **Metodesnitazene** | | **Etodesnitazene** | | **Prodestonitazene** | |
| | SP1 | SP3 | SP1 | SP3 | SP1 | SP3 | SP1 | SP3 |
| Glide | -7.4 | — | — | -6.3 | -7.0 | -6.2 | — | -4.4 |
| MOE | -9.1 | -8.3 | -8.1 | -7.2 | -8.5 | -8.6 | -8.5 | -8.3 |
| Autodock | -7.9 | -7.2 | -7.5 | — | -7.7 | — | -8.0 | — |

Scores from Autodock and from MOE are in kcal/mol, while scores from Glide are unitless. Poses are defined by the group that occupies sp1: nitro (sp1 pose in manuscript) or oxy (flipped pose). Note, only poses that show a potential salt bridge between the piperidine and Asp147 are retained. A hydrogen bond constraint was added during docking in both MOE and Glide; for Autodock a similar constraint was added after docking by imposing a distance cutoff of 4 Å between the piperidine nitrogen and the nearest carboxylate oxygen of Asp147, and ensuring the hydrogen on the piperidine nitrogen was closer to the carboxylate oxygen than the piperidine nitrogen.

Table S4: Docking poses and scores of 8 nitazenes based on the cryo-EM structure model of $\mu$OR:MP complex.

| | Metonitazene | | Etonitazene | | Protonitazene | | Isotonitazene | |
|---|---|---|---|---|---|---|---|---|
| | SP2 | SP3 | SP2 | SP3 | SP2 | SP3 | SP2 | SP3 |
| Glide | -6.2 | — | -6.5 | — | -6.7 | — | -6.5 | -4.1 |
| MOE | -6.2 | -7.1 | -8.1 | -6.6 | -8.0 | -7.0 | -6.8 | -6.9 |
| Autodock | -6.9 | — | -6.9 | — | -6.4 | — | -7.2 | -6.6 |
| | **Butonitazene** | | **Metodesnitazene** | | **Etodesnitazene** | | **Prodestonitazene** | |
| | SP2 | SP3 | SP2 | SP3 | SP2 | SP3 | SP2 | SP3 |
| Glide | -6.2 | — | -6.3 | — | -6.2 | -5.7 | -5.4 | -5.6 |
| MOE | -7.2 | -7.0 | -6.7 | -6.3 | -6.8 | -6.7 | — | -6.4 |
| Autodock | -6.5 | — | -6.0 | -5.9 | -6.4 | -6.8 | -6.7 | -6.5 |

Scores from Autodock and from MOE are in kcal/mol, while scores from Glide are unitless. Poses are defined by the group that occupies sp2: nitro (sp2 pose in manuscript) or oxy (flipped pose). Note, only poses that show a potential salt bridge between the piperidine and Asp147 are retained. A hydrogen bond constraint was added during docking in both MOE and Glide; for Autodock a similar constraint was added after docking by imposing a distance cutoff of 4 Å between the piperidine nitrogen and the nearest carboxylate oxygen of Asp147, and ensuring the hydrogen on the piperidine nitrogen was closer to the carboxylate oxygen than the piperidine nitrogen.

Table S5: Occupancy of the water-mediated hydrogen bonds between the nitro-containing nitazenes and polar residues of $\mu$OR listed in Figure 7.

| | | SP2 | | SP3 | |
|---|---|---|---|---|---|
| | Q124 | Y75 | H319 | W318 | H297 |
| Meto- | 15% | 18% | — | 6% | — |
| Eto- | — | 57% | 9% | 9% | — |
| Proto- | 4% | 30% | 6% | 15% | — |
| Isoto- | — | 39% | 12% | 2% | — |
| Buto- | 17% | 61% | 48% | 15% | — |

The donor-acceptor cutoff distance was set to 3.5 Å, while the donor-hydrogen-acceptor angle cutoff was set to 120 degrees.

Table S6: A complete list of receptor-ligand contacts and occupancies from the cMD refinement simulations

| | TM1 | TM2 | | | | | | |
|---|---|---|---|---|---|---|---|---|
| | Y75 | A117 | L121 | Q124 | S125 | N127 | Y128 | W133 |
| meton | 0.92 | 0.96 | 0.92 | 1.00 | 0.51 | 0.05 | 0.69 | 0.00 |
| etoni | 1.00 | 0.29 | 0.99 | 1.00 | 0.97 | 0.03 | 1.00 | 0.00 |
| proto | 0.99 | 0.59 | 0.69 | 0.99 | 0.60 | 0.04 | 0.97 | 0.00 |
| buton | 0.91 | 0.00 | 0.03 | 0.86 | 0.00 | 0.00 | 1.00 | 0.00 |
| isoto | 1.00 | 0.73 | 1.00 | 1.00 | 0.98 | 0.03 | 1.00 | 0.00 |

| | TM3 | | | | | | |
|---|---|---|---|---|---|---|---|
| | V143 | I144 | D147 | Y148 | N150 | M151 | F152 |
| meton | 0.00 | 0.01 | 1.00 | 1.00 | 0.78 | 1.00 | 0.02 |
| etoni | 0.01 | 0.19 | 1.00 | 1.00 | 0.04 | 1.00 | 0.07 |
| proto | 0.00 | 0.05 | 1.00 | 0.65 | 0.06 | 0.96 | 0.02 |
| buton | 0.00 | 0.01 | 1.00 | 1.00 | 0.00 | 1.00 | 1.00 |
| isoto | 0.00 | 0.18 | 1.00 | 0.80 | 0.07 | 0.99 | 0.00 |

| | TM5 | | | | | |
|---|---|---|---|---|---|---|
| | E229 | L232 | K233 | V236 | F237 | A240 |
| meton | 0.0 | 0.7 | 1.0 | 1.0 | 0.4 | 0.0 |
| etoni | 0.0 | 0.3 | 0.9 | 1.0 | 0.6 | 0.0 |
| proto | 0.0 | 0.2 | 0.9 | 1.0 | 0.8 | 0.2 |
| buton | 1.0 | 1.0 | 1.0 | 1.0 | 0.0 | 0.6 |
| isoto | 0.0 | 0.4 | 0.9 | 1.0 | 0.4 | 0.2 |

| | TM6 | | | | |
|---|---|---|---|---|---|
| | W293 | I296 | H297 | V300 | I301 |
| meton | 0.5 | 1.0 | 0.8 | 1.0 | 0.3 |
| etoni | 1.0 | 1.0 | 1.0 | 1.0 | 0.0 |
| proto | 0.1 | 1.0 | 0.9 | 1.0 | 0.6 |
| buton | 1.0 | 1.0 | 0.4 | 1.0 | 0.0 |
| isoto | 1.0 | 1.0 | 0.8 | 1.0 | 0.0 |

| | TM7 | | | | |
|---|---|---|---|---|---|
| | W318 | H319 | I322 | G325 | Y326 |
| meton | 1.0 | 0.2 | 1.0 | 0.0 | 1.0 |
| etoni | 1.0 | 0.9 | 1.0 | 0.9 | 1.0 |
| proto | 1.0 | 0.6 | 1.0 | 0.8 | 1.0 |
| buton | 1.0 | 1.0 | 1.0 | 0.0 | 1.0 |
| isoto | 1.0 | 1.0 | 1.0 | 0.3 | 1.0 |

| Header column | |
|---|---|
| (red) | Fentanyl (8EF5) |
| (blue) | MP (7T2G) |
| (green) | Both |
| (white) | Neither |

Distance cutoff: 4.5 angstrom

Coloring is based on

### Supplemental figures

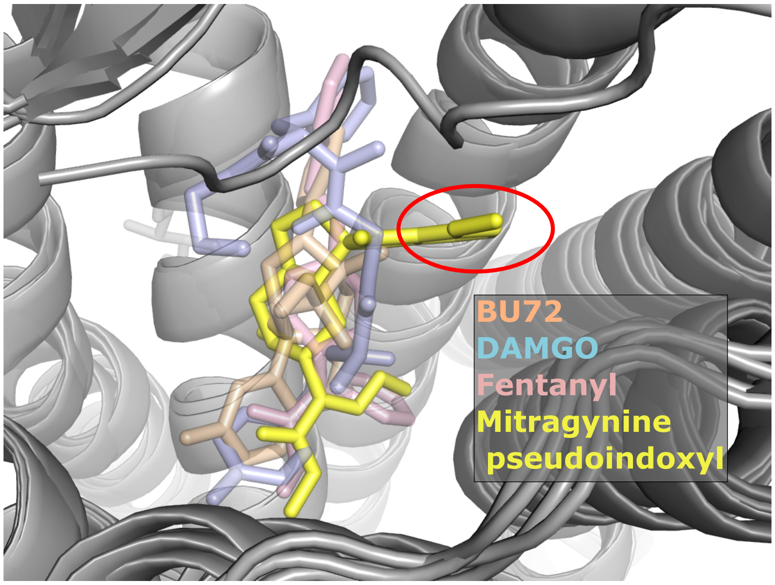


Figure S1: **Mitragynine pseudoindoxyl occupies a unique region compared to other agonists.** Comparison of the resolved pose of four agonists: BU72 (PDB: 5C1M), DAMGO (PDB:8EFQ), fentanyl (PDB:8EF5), and mitragynine pseudoindoxyl (PDB: 7T2G). Instead of occupying the hydrophobic space between TM2 and TM3 (SP1), mitragynine pseudoindoxyl positions an indole on the opposing side of TM2, between TM2, TM7, and TM1 (SP2).

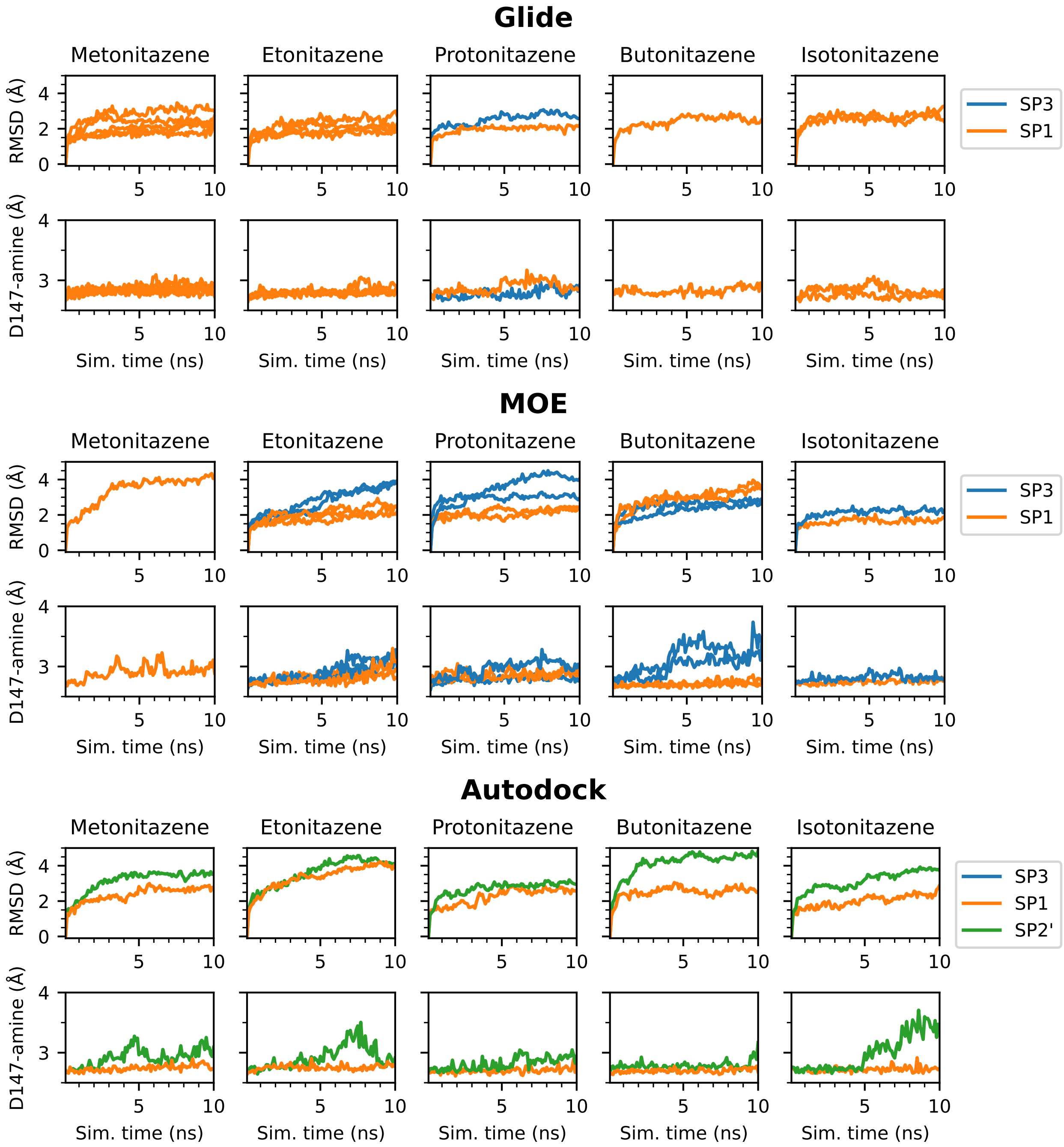


Figure S2: **Stabilities of the binding poses of the nitro-containing nitazenes in $\mu$OR assessed by the binding pose metadynamics.** For each software, the top panel shows the time series of the average ligand RMSD, and the bottom panel shows the time series of the average distance between the triethylamine nitrogen and the nearest carboxylate oxygen of Asp147 for the different poses. Nitazenes containing a nitro group consistently show SP1 being most stable with the lowest average RMSD after 10 ns and with no disruption of the amine-Asp147 salt bridge. Each pose was predicted using the BU72:$\mu$OR complex (PDB: 5C1M) as a docking template.

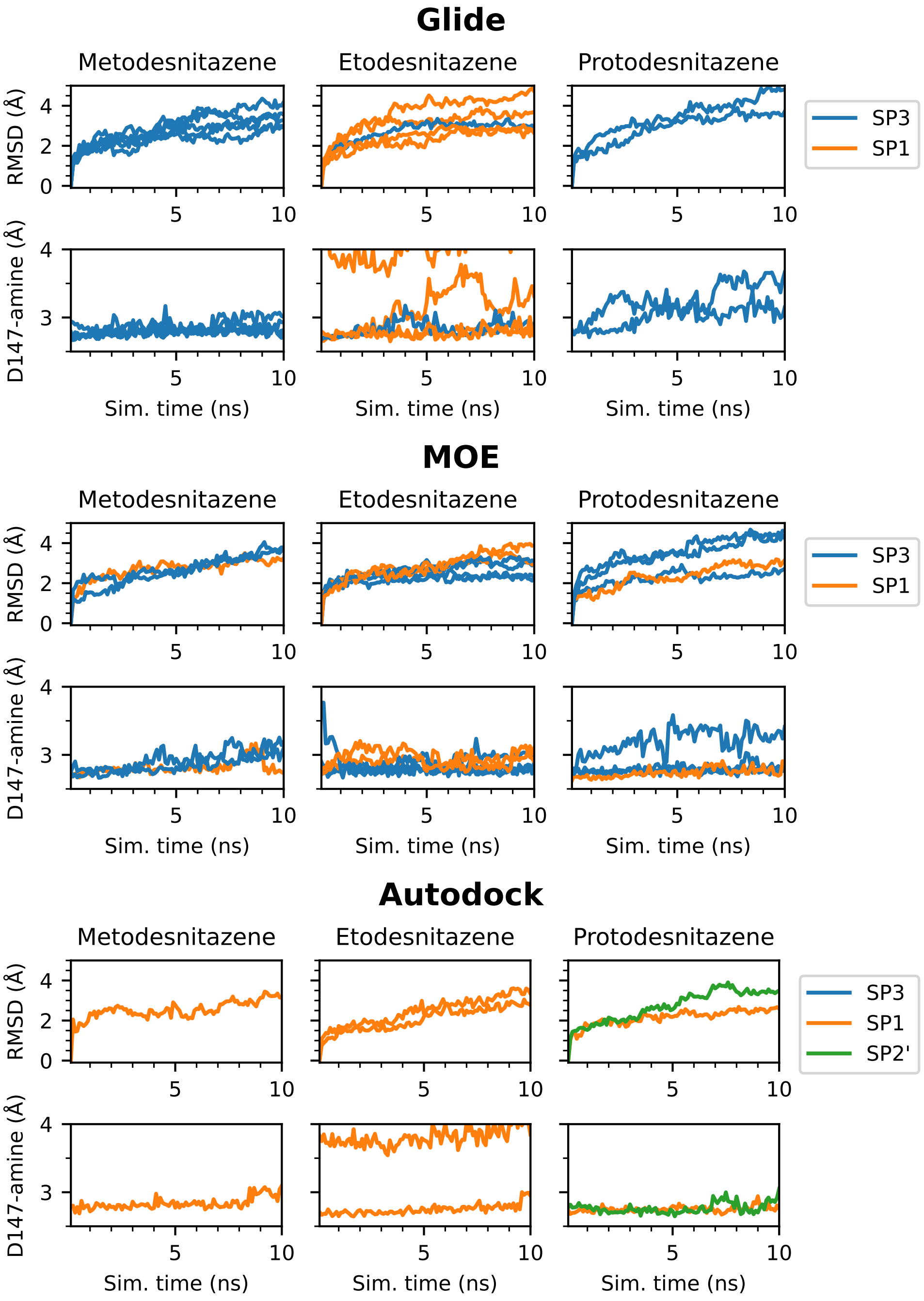


Figure S3: **Stabilities of the binding poses of the nitro-less nitazenes in $\mu$OR assessed by the binding pose metadynamics.** For each software, the top panel shows the time series of the average ligand RMSD, and the bottom panel shows the time series of the average distance between the triethylamine nitrogen and the nearest carboxylate oxygen of Asp147 for the different poses. Metodes-, etodes-, protodesnitazenes do not show a consistent trend. Autodock was the only docking software to produce poses placing the nitro group towards SP2 (SP2' pose). Each pose was predicted using the BU72:$\mu$OR complex (PDB: 5C1M) as a docking template.

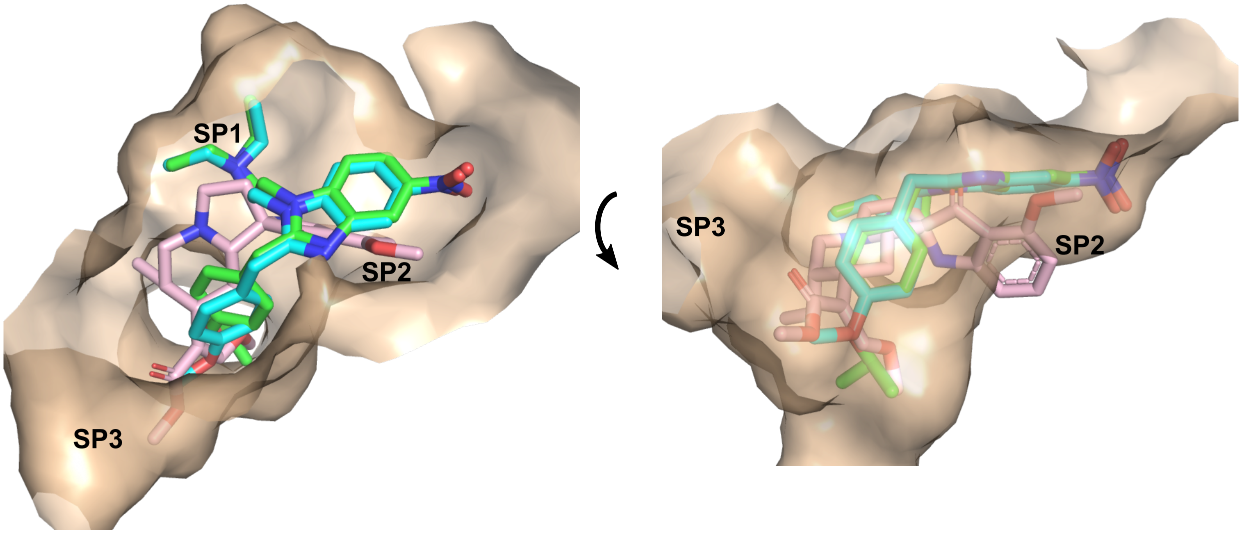


Figure S4: **Autodock predicts a variation of the SP2 pose (SP2') when docking select nitazenes to the BU72:$\mu$OR complex.** Visualization of the SP2' pose predicted by Autodock for metonitazene (cyan) and isotonitazene (green); for reference the MP:$\mu$OR complex (PDB: 7T2G) was superimposed and shown in pink. Autodock placed the nitro group above the SP2 pocket where the indole of MP resides, as Gln124 (not shown for clarity) occupies SP2. Meanwhile the *N*-diethyl occupies SP1 (instead of the nitro group or oxyl tail), and the oxyl tail extends towards the center of the receptor–leaving SP3 empty.

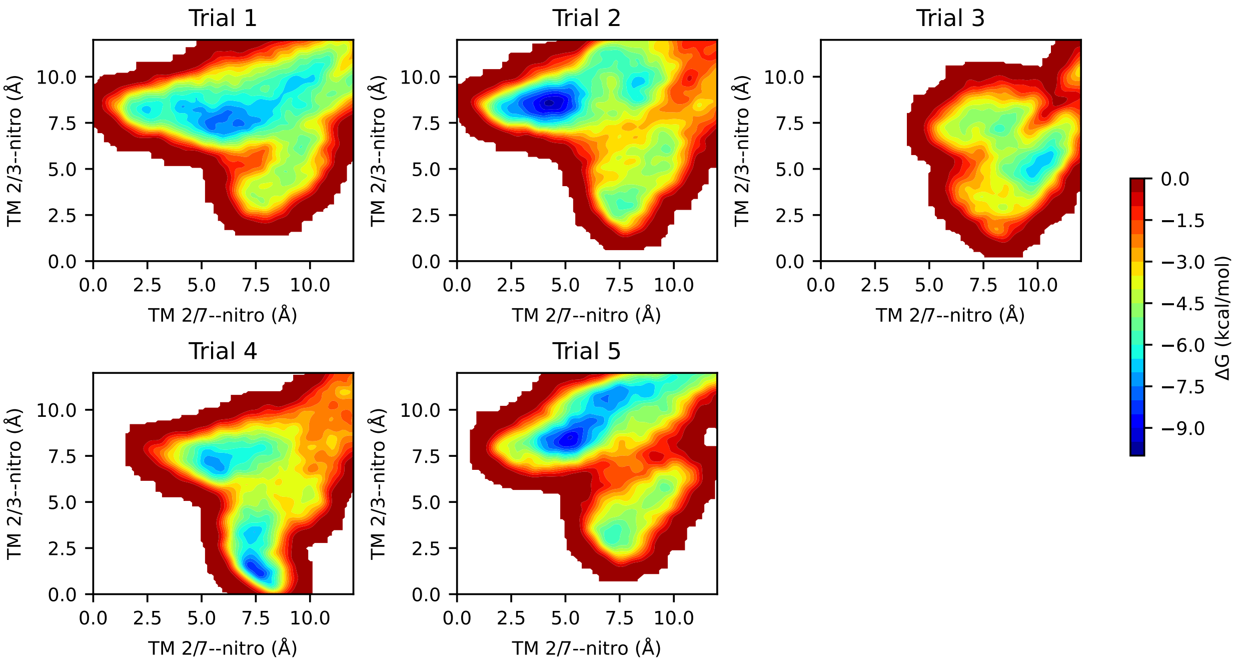


Figure S5: **Metadynamics simulations estimate the SP2 pose is more favorable than the SP3 pose.** Approximate free energy landscape from five independent metadynamics simulations. Of the five simulations, two (trials 3 and 4) did not sample the SP2 binding mode.

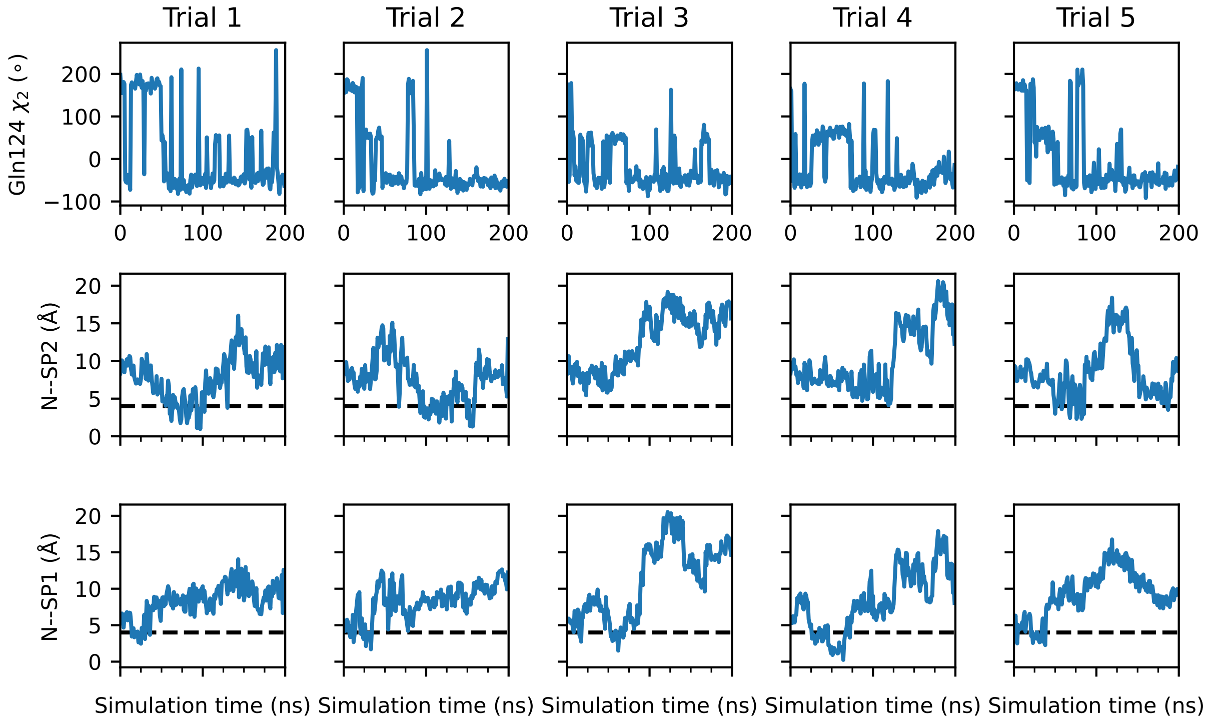


Figure S6: **Gln124 changes conformation once the nitro group of etonitazene exits SP1.** Time evolution of the $\chi_2$ of Gln124 (top row), as well as the distance from the nitro nitrogen of etonitazene to the center of mass of SP2 (middle row) and SP1 (bottom row), for each of the metadynamics trajectories starting from the SP1-binding pose. The black dashed line at 4 Å indicates the distance at which the nitro group is positioned at either SP2 or SP1.

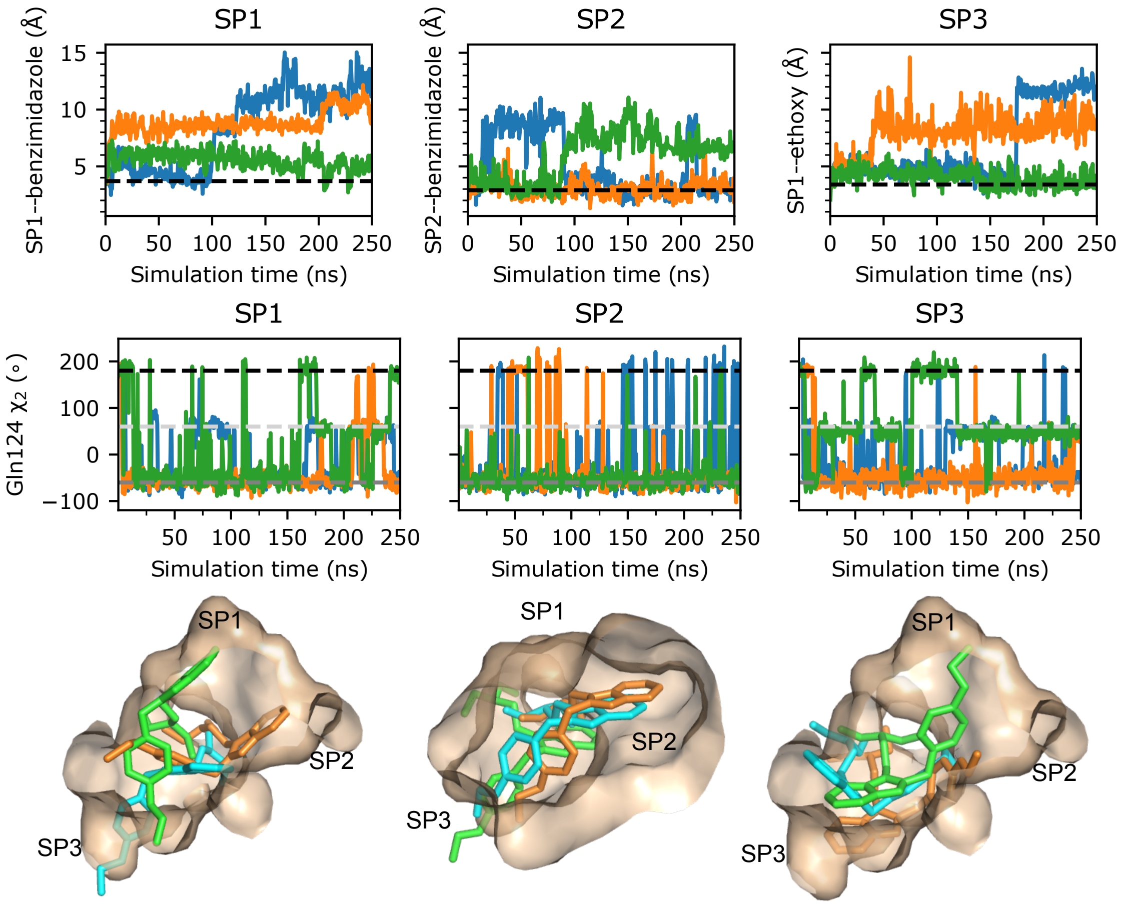


Figure S7: **cMD simulations suggest that SP2 is more stable than SP1 and SP3 poses for etodesnitazene.** Results from cMD simulations starting from the top SP1, SP2, or SP3 docked poses of etodesnitazene. Three independent simulations (blue, orange, and green) were conducted for each pose. **First row.** Distance between the benzimidazole of etodesnitazene and the occupied subpocket as a function of time for SP1, SP2, and SP3 binding poses. The initial pocket distance for each pose is shown as a black dashed line. **Second row.** $\chi_2$ of Gln124 as a function of time for each simulation of etodesnitazene. **Third row.** RMSD of etodesnitazene, with respect to the initial pose, as a function of time for each simulation. **Fourth row.** Visualization of etodesnitazene within the binding pocket for the final frame of each replica.

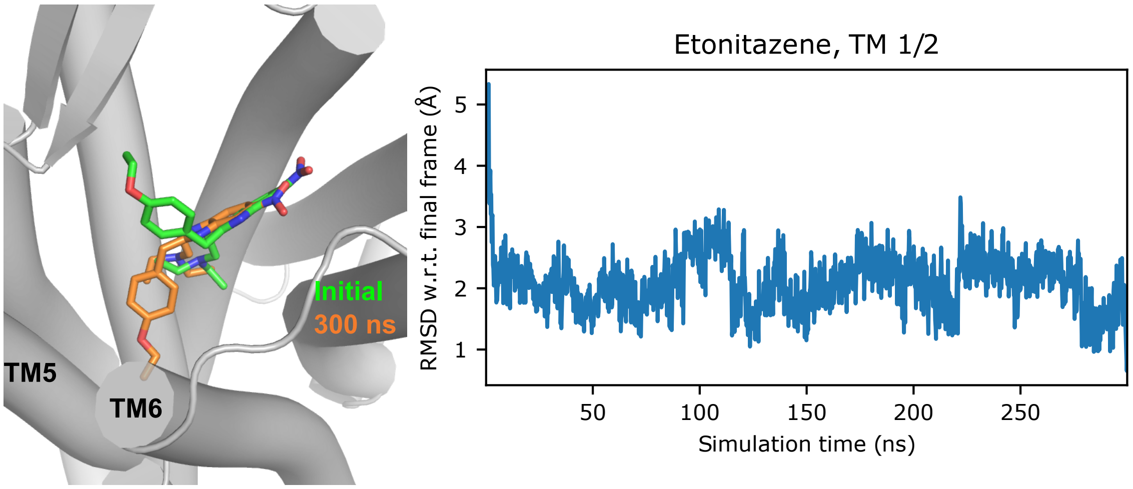


Figure S8: **Alkoxy tail of etonitazene is placed into SP3 after cMD refinement. Left.** Visualization of the top SP2 docked pose of etonitazene (from Glide; based on $\mu$OR:MP template; green) and the final frame of a 300 ns cMD simulation (orange). **Left.** RMSD of etonitazene with respect to the final frame of the simulation as a function of time.

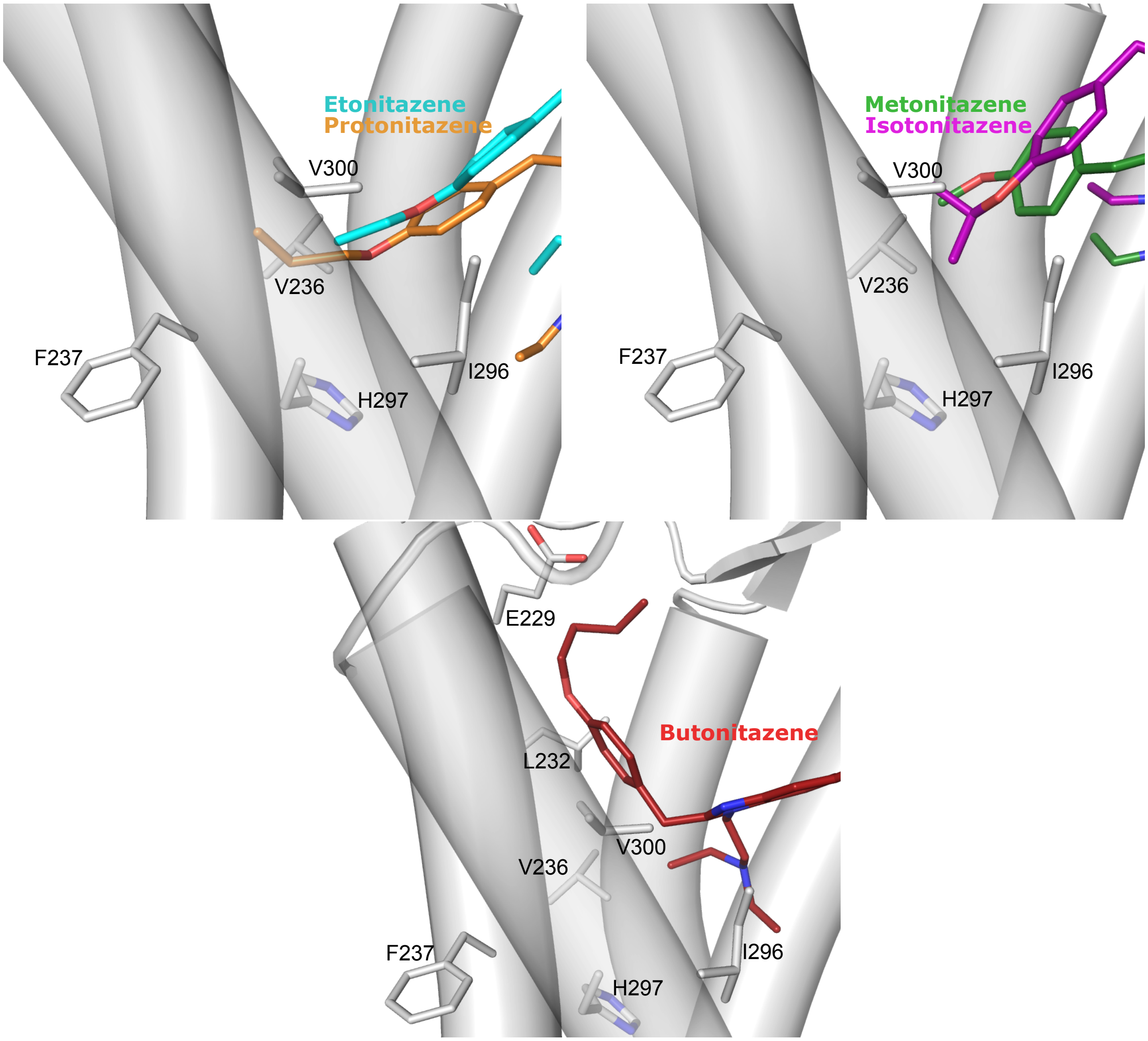


Figure S9: **Visualization of the alkoxy tails of the five nitro-containing nitazenes in SP3. Top left.** Interactions of eto- and protonitazenes with SP3 residues. **Top right.** Interactions of meto- and isotonitazenes with SP3 residues. **Bottom.** The butoxy tail of butonitazene is shifted upwards. Although it maintains the interactions with the residues flanking SP3, it loses contact with the backbone of F237 and gains contacts with Glu229 and Leu232 above SP3. A complete list of contacts and their occupancies is given in Table S6. The final frames of the 100-ns cMD refinement simulations of the SP2 pose for meto-, proto, isoto-, and butonitazenes are used. For etonitazene, the final frame of the 300-ns cMD refinement simulation of the SP2 pose is used.

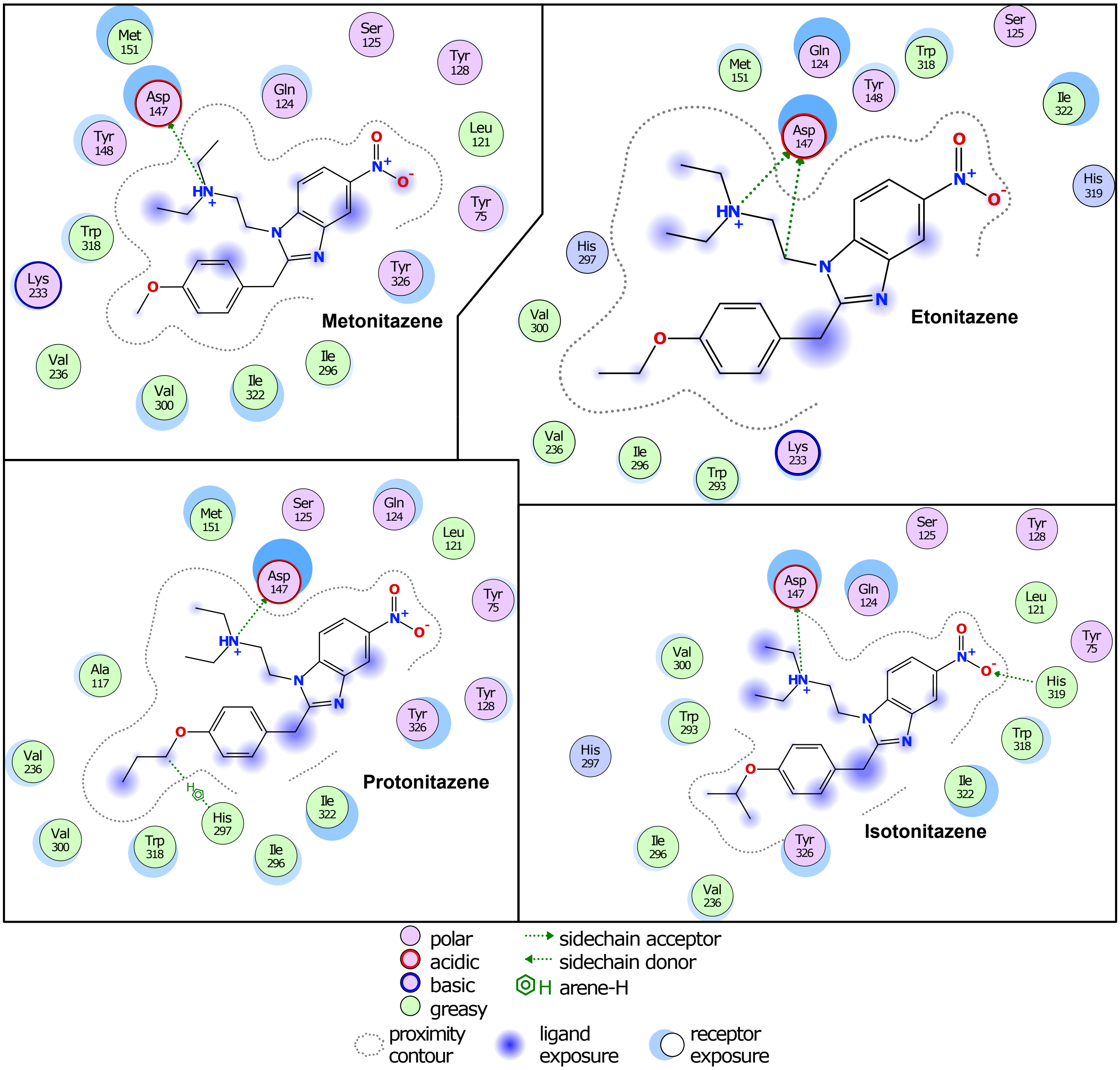


Figure S10: **Interactions between four nitro-containing nitazenes and $\mu$-opioid receptor.** 2D visualization of receptor-ligand interactions for meto-, eto-, proto-, and isotonitazenes. The final frame of a 100 ns simulation was used. Visualization and interaction analysis was performed using MOE.

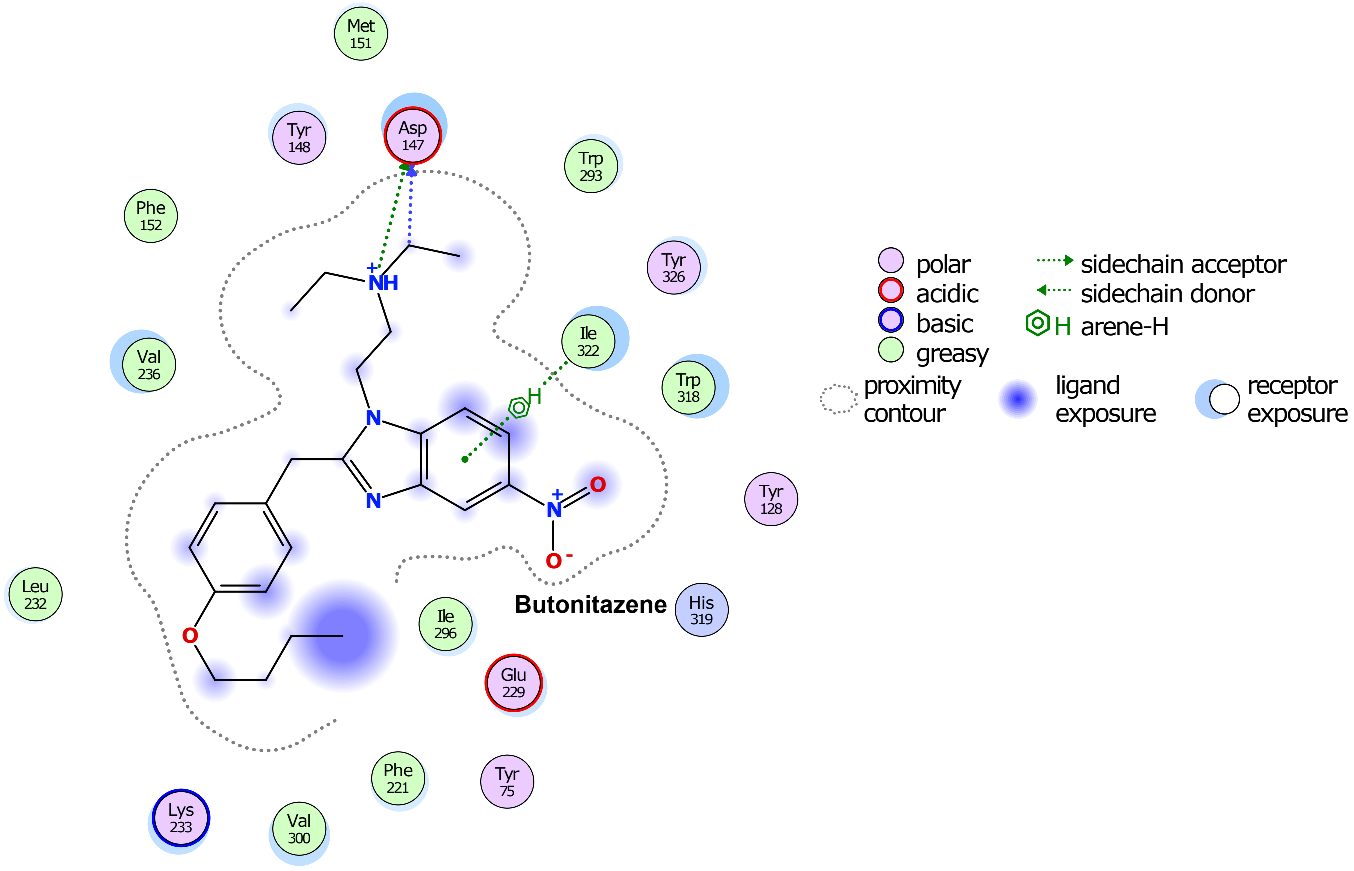


Figure S11: **Interactions between butonitazene and $\mu$-opioid receptor.** 2D visualization of receptor-ligand interactions. The final frame of a 100 ns simulation was used. Visualization and interaction analysis was performed using MOE.
